# Identification of chemical features that influence mycomembrane permeation and antitubercular activity

**DOI:** 10.1101/2025.02.27.640664

**Authors:** Irene Lepori, Zichen Liu, Nelson Evbarunegbe, Shasha Feng, Turner P. Brown, Kishor Mane, Shivangi, Mitchell Wong, Amir George, Taijie Guo, Jiajia Dong, Joel S. Freundlich, Wonpil Im, Anna G. Green, Marcos M. Pires, M. Sloan Siegrist

## Abstract

Tuberculosis (TB), caused by *Mycobacterium tuberculosis* (Mtb), is the deadliest single-agent infection worldwide. Current antibiotic treatment for TB takes a minimum of four months, underscoring the need for better therapeutics. The unique mycobacterial cell envelope, particularly the outermost mycomembrane, has long been thought to promote intrinsic antibiotic resistance by limiting compound entry into Mtb. Understanding chemical features that influence permeation across the mycomembrane may enable more accurate predictions of whole cell anti-Mtb activity, leading to development of more effective antibacterials. Here we query the mycomembrane permeation of over 1500 azide-tagged compounds in live Mtb with the bioorthogonal click chemistry-based assay PAC-MAN. We use cheminformatics and machine learning to identify chemical features associated with mycomembrane permeation and show that they have predictive value via systematic modification of two small molecule series. Additionally, we find that chemical features that influence mycomembrane permeation correlate with anti-Mtb activity in large compound libraries. These findings suggest that the mycomembrane is indeed a significant barrier to whole cell activity in Mtb and provide a rational framework for designing or modifying compounds to overcome this barrier.

## Introduction

Tuberculosis (TB) caused by *Mycobacterium tuberculosis* (Mtb) consistently claims over one million lives each year^1^. Antibiotic treatment is lengthy, complex, and variably effective. High doses are often required for TB treatment, which in turn increase the risk of side effects and reduce patient compliance. Better drugs and drug regimens to target TB are urgent public health priorities.

An outstanding challenge in developing or improving drugs for TB, as with many other diseases, is ensuring that drug candidates accumulate within the cells implicated in disease pathogenesis^2, 3^. To achieve whole cell activity, a molecule must overcome several barriers, including membranes, efflux, and metabolism, to accumulate with the kinetics and at the concentrations required to effectively engage its target. The physicochemical features or moieties of a molecule that enable it to overcome individual accumulation barriers are divergent and sometimes at odds^4^.

Moreover, even within bacteria there are likely inter-species and genera differences in molecule accumulation. For example, the extent and/or pattern of accumulation for certain antibacterials, nutrients, and other molecules differ in mycobacteria relative to Gram-negative bacteria^5-8^. This inference has been generalized with the determination of molecular correlates of accumulation for the Gram-negative species *Escherichia coli* and *Pseudomonas aeruginosa*^9, 10^ and the mycobacterial pathogen *Mycobacterium abscessus*^11^. Strikingly, physicochemical properties alone, including those associated with *E. coli* or *P. aeruginosa* accumulation, fail to predict *M. abscessus* accumulation and necessitated a deep learning approach to identify correlates^11^. These findings indicate that molecule accumulation in Mtb is likely to be a complex phenotype.

Identifying chemical features that promote Mtb cell accumulation may enable more rapid development of new tuberculosis treatments. One way to address the complexity of accumulation is to focus on chemical features that enable a molecule to overcome a single barrier. For example, it has long been assumed that Mtb and other mycobacterial pathogens are intrinsically resistant to certain drugs in part because of the impermeability of their outermost mycomembrane^6, 12, 13^. This assumption is predicated on studies showing that mycomembrane disruptions sensitize mycobacteria to a subset of antibacterials^6, 7, 12-15^. However, activity is an indirect proxy for uptake; collateral metabolic dysfunction from cell envelope perturbation^16-19^ may also impact drug sensitivity.

Here we addressed the complexity of Mtb accumulation by first screening a small molecule library for mycomembrane permeation and then deploying machine learning to discover chemical features that influence this phenotype. We used our recently-developed, click chemistry-based assay <u>P</u>eptidoglycan <u>A</u>ccessibility <u>C</u>lick-<u>M</u>ediated <u>A</u>ssessme<u>N</u>t (PAC-MAN^7, 20^; **Fig. 1a**) to measure the permeation of over 1500 small molecules across the mycomembrane of live Mtb. Cheminformatics analyses suggest that the rules for mycomembrane permeability are not simple, as physicochemical properties have scaffold-dependent effects. To capture structure-function relationships more holistically, we trained a neural network to predict mycomembrane permeability for any compound, then queried for chemical features predictive of permeability. *De novo* chemical synthesis and mycomembrane permeation testing of a two small molecule series confirmed that the chemical features identified by cheminformatics and machine learning are predictive. Finally, we showed that scaffolds and other chemical features that predict mycomembrane permeation also correlate with anti-Mtb activity. Our work supports the long-standing hypothesis that the mycomembrane is a major barrier to whole cell activity in Mtb and suggests that improving the ability of a molecule to permeate the mycomembrane may aid the rational design or redesign of more effective drugs for tuberculosis.

**Figure 1.**
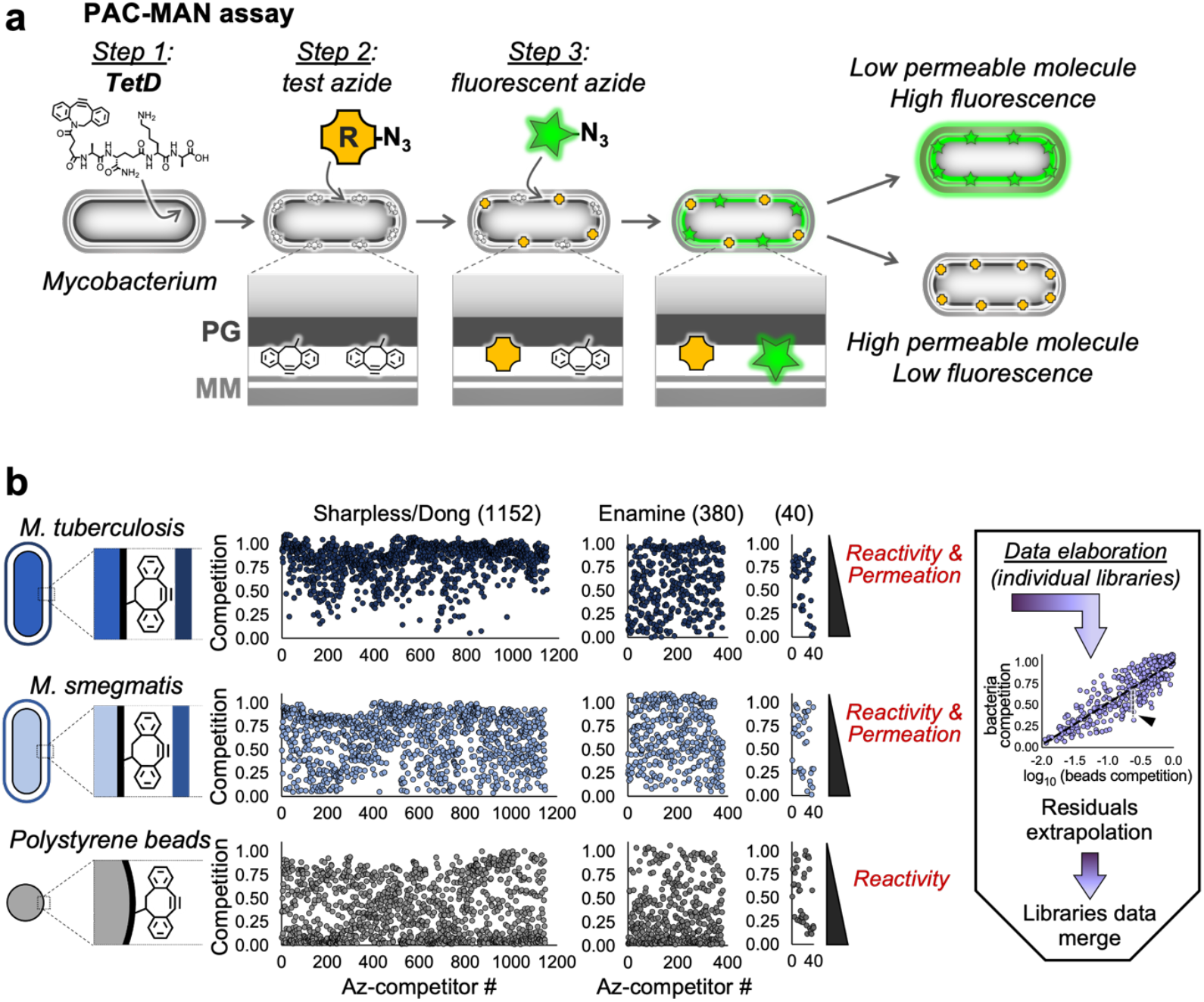
High-throughput screening of mycomembrane permeability. (**a**) Schematic of PAC-MAN assay (adapted from^7^). In *Step 1*, Mtb or Msm is incubated with the DBCO-bearing **TetD** probe to tag cell wall peptidoglycan. Mycobacteria are then exposed to an azide-modified test molecule (*Step 2*) followed by an azide fluorophore (*Step 3*). Test azides that do not permeate the mycomembrane do not access cell wall-embedded DBCO. DBCO that do not react in *Step 2* are free to react with azide fluorophores in *Step 3*, resulting in high fluorescence. Conversely, test azides that permeate the mycomembrane can access and react with DBCO in *Step 2*, preventing DBCO from reacting with azide fluorophores in *Step 3* and resulting in low fluorescence. (**b**) PAC-MAN screening of three test azide libraries: Sharpless/Dong (1152), Enamine (380), or other commercial sources (40). Screening was performed on Mtb, Msm, and DBCO-polystyrene beads to normalize for test azide reactivity. Standardized residuals were extracted from log-linear regression analyses for individual libraries as shown. Values for standardized residuals are inversely proportional to mycomembrane permeability of test azide.

## Results

### High-throughput screening of mycomembrane permeability

The PAC-MAN assay (**Fig. 1a**) first proceeds via metabolic labeling of peptidoglycan, the cell wall polymer immediately beneath the arabinogalactan-mycomembrane layer of the cell envelope, with a dibenzocyclooctyne (DBCO) probe. A portion of the peptidoglycan-embedded DBCO is captured via bioorthogonal, strain-promoted alkyne-azide cycloaddition (SPAAC^21-23^) with azide-tagged test molecules. The remaining DBCO groups are revealed via SPAAC with a fluorescent azide. The level of mycobacterial cell fluorescence, measured by flow cytometry, inversely correlates with permeation of the azide test molecule across the mycomembrane^7, 20^. We used PAC-MAN to screen Mtb mc^2^6206 (H37Rv Δ*panCD* Δ*leuCD*)^24^ and the model organism *Mycobacterium smegmatis* mc^2^155 (Msm)^25^ with 1572 azide test molecules from three sources: 1152 synthesized via fluorosulfuryl azide chemistry from the corresponding primary amine (Sharpless/Dong^26^), 380 purchased from Enamine, and 40 purchased from various commercial sources (**Fig. 1b**; **Fig. S1**).

The latter two sources contain molecules with primary amines. To control for the intrinsic reactivities of azide test molecules toward DBCO^7, 20^, we also screened DBCO-functionalized polystyrene beads that have bacteria-like dimensions but are devoid of a permeability barrier (**Fig. 1b**). Previously we controlled for reactivity and calculated mycomembrane permeation by exposing DBCO-labeled beads and mycobacteria to different concentrations of azide test molecules, then calculating the difference in concentration of test azide molecule that competes fluorescence by 50% in the two systems (Δlog_10_CC_50_)^7^. For higher-throughput calculation of mycomembrane permeation, we instead exposed DBCO-labeled beads and mycobacteria to fixed concentrations of test azide molecules then performed log-linear regression analyses to calculate the differences between observed and expected fluorescence (standardized residuals; **Fig. 1b**; **Fig. S2**). The standardized residuals for compounds shared between the Sharpless/Dong and Enamine libraries were consistent (**Fig. S3**).

### Cheminformatics analyses identify chemical features associated with mycomembrane permeability

We first analyzed the correlation between the presence of small chemical scaffolds and mycomembrane permeability. We used a permissive strategy (**Fig. 2a, Fig. S4**), which permits the scaffold of interest to be fused or not with other rings, as well as a “greedy” strategy that excludes the ring-fused versions of the scaffold (**Fig. 2b**). For both Msm and Mtb we found that aromatic nitrogen-containing scaffolds like indole, imidazole, or pyrazole correlate positively with permeation (negative medians for standardized residuals), while scaffolds like cyclopentane or cyclohexane correlate negatively (positive medians).

**Figure 2.**
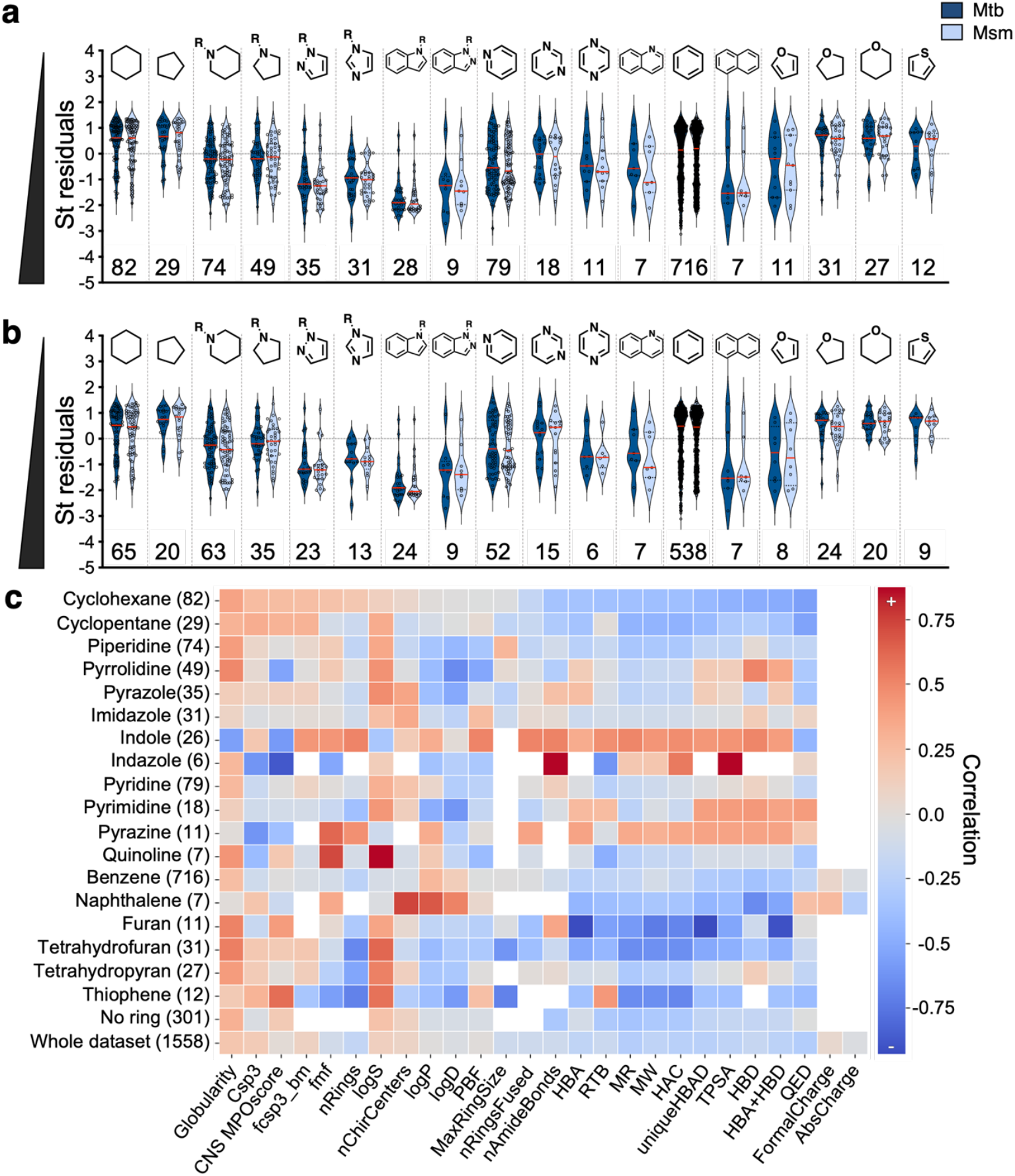
Chemical scaffolds and physicochemical properties that correlate with mycomembrane permeability. (**a**) Mycomembrane permeability (standardized [St] residuals) for test azides grouped by the presence of a specific ring-containing scaffold, indicated on top. (**b**) “Greedy” scaffold analysis excludes the fused rings structures. Numbers below the plots are the number of compounds in each group. (**c**) Correlation between mycomembrane permeability and selected physicochemical properties for all test azides (whole dataset, *bottom*) and for subgroups bearing the scaffolds plotted in (**a-b**). Numbers in brackets are the number of compounds in each group. Red and blue respectively denote positive and negative correlations between the indicated physicochemical properties and mycomembrane permeation. For descriptors explanation see **Figure S1**.

We next investigated the correlation between mycomembrane permeation and physicochemical properties (**Fig. 2c**). When analyzed as a whole, our Mtb dataset did not reveal obvious correlations. Notably, the physicochemical properties previously shown to correlate with Gram-negative accumulation^9, 10, 27^ were not associated with mycomembrane permeation (**Fig. S5**). Reasoning that the contributions of physicochemical properties may depend on the structural context in which they occur, we repeated the cheminformatics analyses after grouping compounds by the presence of specific scaffolds as in **Fig. 2a**. We found that the effects of physicochemical properties on mycomembrane permeability varied by scaffold. For example, topological polar surface area (TPSA) and log partition coefficient (logP; Crippen logP calculation from RDKit^28^) have strong, positive correlations with mycomembrane permeation when a molecule contains an indazole or naphthalene, respectively, but have weak and/or negative correlations with mycomembrane permeation in the context of most other scaffolds. These observations are significant as lipophilicity is generally viewed as a positive attribute for antitubercular drugs^29^ and underscores the challenge of identifying molecular correlates of Mtb accumulation even for a single barrier.

### Machine learning predicts chemical features that are associated with mycomembrane permeation

We built a machine learning (ML) model, called Mycobacterial Permeability neural Network (MycoPermeNet), to capture the complex relationship between chemical structure and mycomembrane permeability (**Fig. 3a, Fig. S6**). Inspired by recent successes in deep learning for antibacterial discovery^30, 31^, the model takes SMILES strings and Mtb screening data (standardized residuals) as inputs, then uses a two-stage process to predict mycomembrane permeability. First, it learns to generate vector representations of chemical compounds, called embeddings, using a message passing neural network on molecular graphs implemented in Chemprop^32^. It then uses a downstream multilayer perceptron to convert embeddings into a permeability prediction (*i*.*e*., predicted standardized residuals). The two stages are trained independently using an 80-10-10 train-validate-test split to optimize hyperparameters and perform model selection. The multilayer perceptron performed the best on the validation set out of several model architectures tested (**Fig. S6**). Our final model achieves R^2^, RMSE, and MAE respectively of 0.74, 0.50, and 0.37 for the train set and 0.72, 0.37, and 0.41 for the held-out test sets (**Fig. 3b**), indicating a strong relationship between measured and predicted mycomembrane permeability. Moreover, we achieve Spearman rank correlation coefficients of 0.85 on the train set and 0.86 on the held-out test set, demonstrating that our model correctly ranks the relative permeability of compounds even better than it predicts their absolute permeability scores.

**Figure 3.**
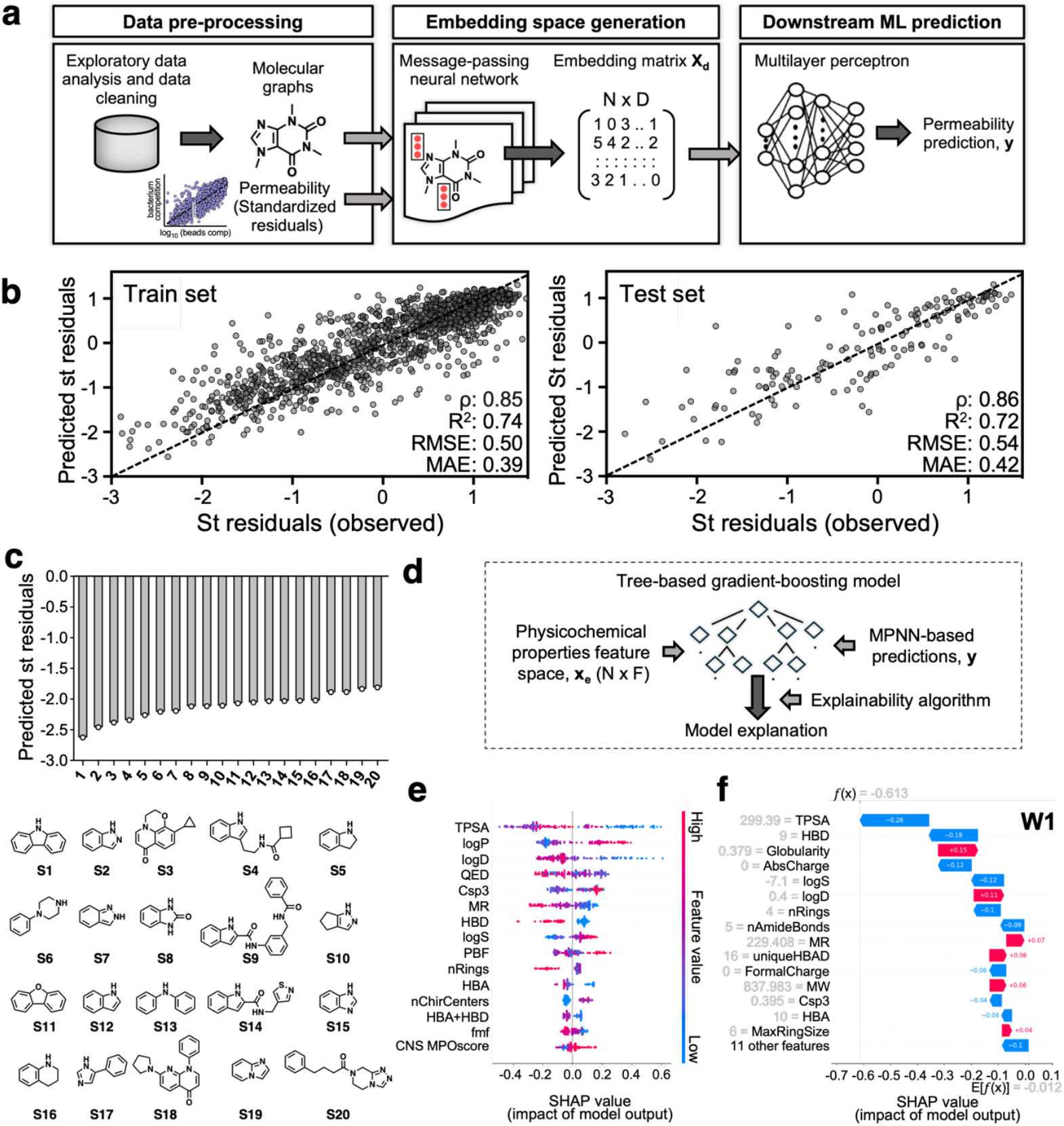
Machine learning model development and interpretation. (**a**) Pipeline for machine learning (ML) model MycoPermeNet development. (**b**) ML-predicted vs. experimentally-observed mycomembrane permeation data (standardized residuals) for both the training (*left*) and validation (*right*) sets. (**c**) The 20 most**-**permeable chemical scaffolds predicted from the ML model. (**d**) Pipeline for surrogate XGBoost ML model SHAP-based interpretability studies. (**e**) Ranking of top-15 (out of 26) physicochemical properties prioritized by the ML model to drive its prediction across all compounds. Molecular descriptors calculated by RDkit and Collaborative Drug Discovery (CDD) Vault. (**f**) Interpretation of the ML predictions for a specific molecule (**W1** peptide, see **Fig. 4a**) from the perspective of physicochemical properties (descriptors). Rationalization and quantification of the positive or negative effect of the top-15 (out of 26) descriptors on molecule permeability across the mycomembrane. For results relative to the full list of descriptors see **Figure S7**. RMSE: squared root of mean squared error; MAE: mean absolute error. For descriptors explanation see **Fig S1**. N = number of compounds; F = number of descriptors. D = dimension of embeddings.

To test whether MycoPermeNet is learning reasonable relationships between molecular structure and mycomembrane permeability, we asked the model to predict permeability scores for every Bemis-Murcko scaffold found in the dataset (n=217). Among the 20 scaffolds predicted as most permeable (**Fig. 3c**) we found various indole or indole-like scaffolds, as well as imidazole- and pyrazole-like scaffolds. The concordance between the ML- and cheminformatics-based (**Fig. 2**) analyses serves as a validation of the ML approach and further reinforces that the presence of certain nitrogen aromatic heterocycles correlates with mycomembrane permeability.

To gain additional insights into the physicochemical properties that correlate with molecule permeation across the mycomembrane we performed interpretability studies of MycoPermeNet (**Fig. 3d-f; Fig. S7**). Because our model is based on input chemical structures, which are meaningful individually but difficult to summarize across our large and diverse dataset, we built a surrogate model^33^ to determine which human-interpretable chemical descriptors have the highest influence on the permeability predictions made by MycoPermeNet. Specifically, we used a surrogate XGBoost^34^ to predict the outputs of MycoPermeNet from 26 hand-selected input features (23 calculated with RDKit^28^ and 3 calculated with Collaborative Drug Discovery [CDD] Vault^35^) that we considered important for mycomembrane permeability. After training our surrogate XGBoost model (R^2^, RMSE, and MAE respectively of 0.84, 0.33, and 0.25 for the train set and 0.83, 0.38, and 0.28 for the held-out test set, where the R^2^ on the held-out test set explains 83 percent of the variance in relationship between the features and MycoPermeNet), we interpret its features using SHAP^36^ (SHapley Additive exPlanations; **Fig. 3d**). In this analysis, the two most influential physicochemical properties are TPSA and logP (**Fig. 3e**; **Fig. S7a**). While SHAP returns properties that drive mycomembrane permeability predictions across the dataset, it can also predict which features are responsible for the permeability prediction for individual compounds (**Fig. 3f**; **Fig. S7b**). This analysis provides both a qualitative and quantitative interpretation of the impact of physicochemical properties on mycomembrane permeability.

### Chemical features that are associated with mycomembrane permeation have high predictive value

We hypothesized that molecular features identified by cheminformatics and/or predicted by MycoPermeNet to correlate with mycomembrane permeability are causative. We first tested this hypothesis with a pentapeptide series (Phe-Lys-Phe-Lys-Phe) in which we systematically substituted phenylalanines for tryptophans (**Fig. 4a**). The side chains of phenylalanine and tryptophan respectively bear benzene, which is weakly and negatively associated with mycomembrane permeability, and indole, which is strongly and positively associated with mycomembrane permeability

**Figure 4.**
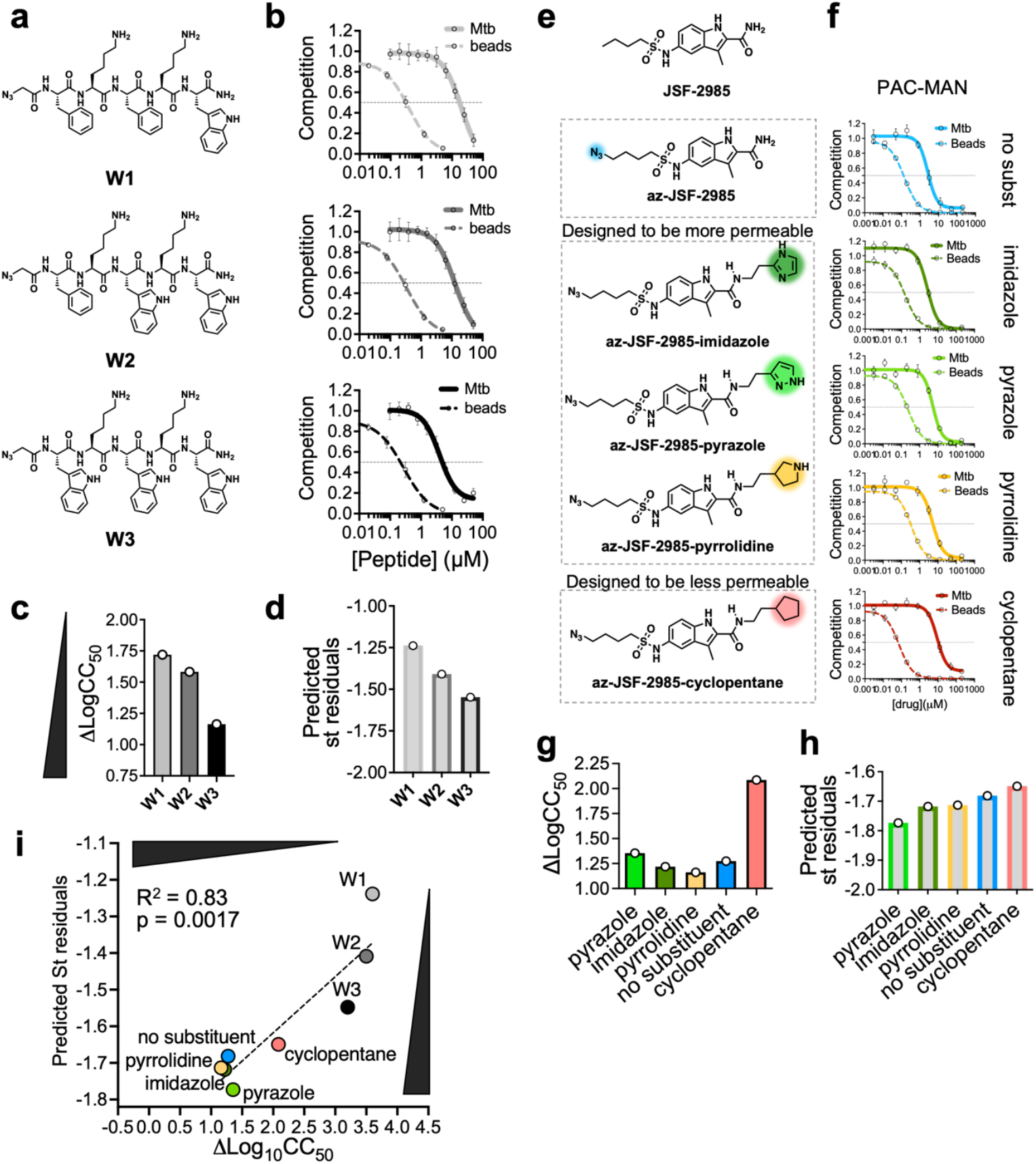
Chemical features identified by cheminformatics and ML influence mycomembrane permeation. (**a-d**) Analysis of a pentapeptide series in which benzene-containing phenylalanines are systematically substituted with indole-containing tryptophans (**W1-3**). The presence of indole or benzene scaffolds are respectively associated with more or less mycomembrane permeability (**Fig. 2a-b**). (**a**) Peptide chemical structures; (**b**) PAC-MAN results for Mtb and beads; (**c**) observed (∆log_10_CC_50_) and (**d**) ML-predicted mycomembrane permeability. (**e-h**) Analysis of an antitubercular candidate series. **JSF-2985** (**e**, *top*) was derivatized to bear an azide and chemical scaffolds that are associated with more (imidazole, pyrrole, pyrrolidine) or less (cyclopentane) mycomembrane permeability (**Fig. 2a-b**). (**f**) PAC-MAN results for Mtb and beads; (**g**) observed (∆log_10_CC_50_) and (**h**) ML-predicted mycomembrane permeability. (**i**) Correlation between observed and ML-predicted mycomembrane permeability across the two compound series.

(**Fig. 2a-b**). We found that substitution of phenylalanines for tryptophans enhanced mycomembrane permeation of the peptides in both Mtb and Msm (**Fig. 4b-c**; **Fig. S8**), consistent with both cheminformatics analyses (**Fig. 2a-b**) and MycoPermeNet predictions (**Fig. 4d**).

We next tested our hypothesis with a small molecule series based on **JSF-2985**, an antitubercular small molecule we previously reported^37^. We synthesized **JSF-2985** analogs via replacement of the 2-position primary amide’s N-H with moieties that correlate positively (imidazole, pyrazole, and pyrrolidine) or negatively (cyclopentane) with mycomembrane permeation (**Fig. 4e**). We chose the 2-position amide N-H as the location of the new substituents based on docking studies that suggest that the amide group is not directly involved in the engagement of the target but instead lies outside the binding pocket and it is thus exposed to the solvent (**Fig. S9**). We found that **JSF-2985** derivatives with imidazole, pyrazole, and pyrrolidine scaffolds permeate the mycomembrane better than the **JSF-2985** derivative bearing cyclopentane (**Fig. 4f-g**), consistent with both cheminformatics analyses (**Fig. 2a-b**) and MycoPermeNet predictions (**Fig. 4h**). The excellent correlation between ML-predicted and observed permeation of the mycomembrane across the two small molecule series (**Fig. 4i**) suggests the potential to rationally tune the ability of a molecule to traverse this barrier.

### Chemical features that predict mycomembrane permeability correlate with whole cell activity

We and many others have shown that mycomembrane disruption sensitizes Mtb to some antibacterials^6, 7, 12-15^. Thus, we hypothesized that mycomembrane permeation is a key determinant of whole cell anti-Mtb activity. Our docking studies (**Fig. S9**) suggested that the **JSF-2985** derivatives have a high probability of target binding. However, we found that Mtb growth inhibition by these molecules did not reflect their mycomembrane permeation profiles (**Fig. S9b-c**), highlighting the complex relationship between molecule accumulation and activity.

We reasoned that a larger dataset, both in terms of the number and size of molecules, would better test our hypothesized relationship between mycomembrane permeation and activity. To this end, we performed a retrospective analysis on a small molecule library previously screened for anti-Mtb activity^38, 39^. We looked for sets of molecules within the ∼200k Molecular Libraries Small Molecule Repository (MLSMR; data obtained from CDD database^35^, Burlingame, CA. www.collaborativedrug.com) collection that have similar structures but differ in the presence or absence of mycomembrane permeability-promoting scaffolds (**Fig. 2a-b**) in peripheral positions. We chose this scheme to decrease the likelihood of the scaffold contributing directly to target engagement. In the four distinct molecule sets that met our criteria (**Fig. 5a-b**; **Fig. S10**), compound activity correlated with the presence of these scaffolds and, more generally, with ML-predicted mycomembrane permeation (note that standardized residuals are inverted in **Fig. 5, Fig. S10**, and **Fig. S12** for more intuitive comparisons to activity).

**Figure 5.**
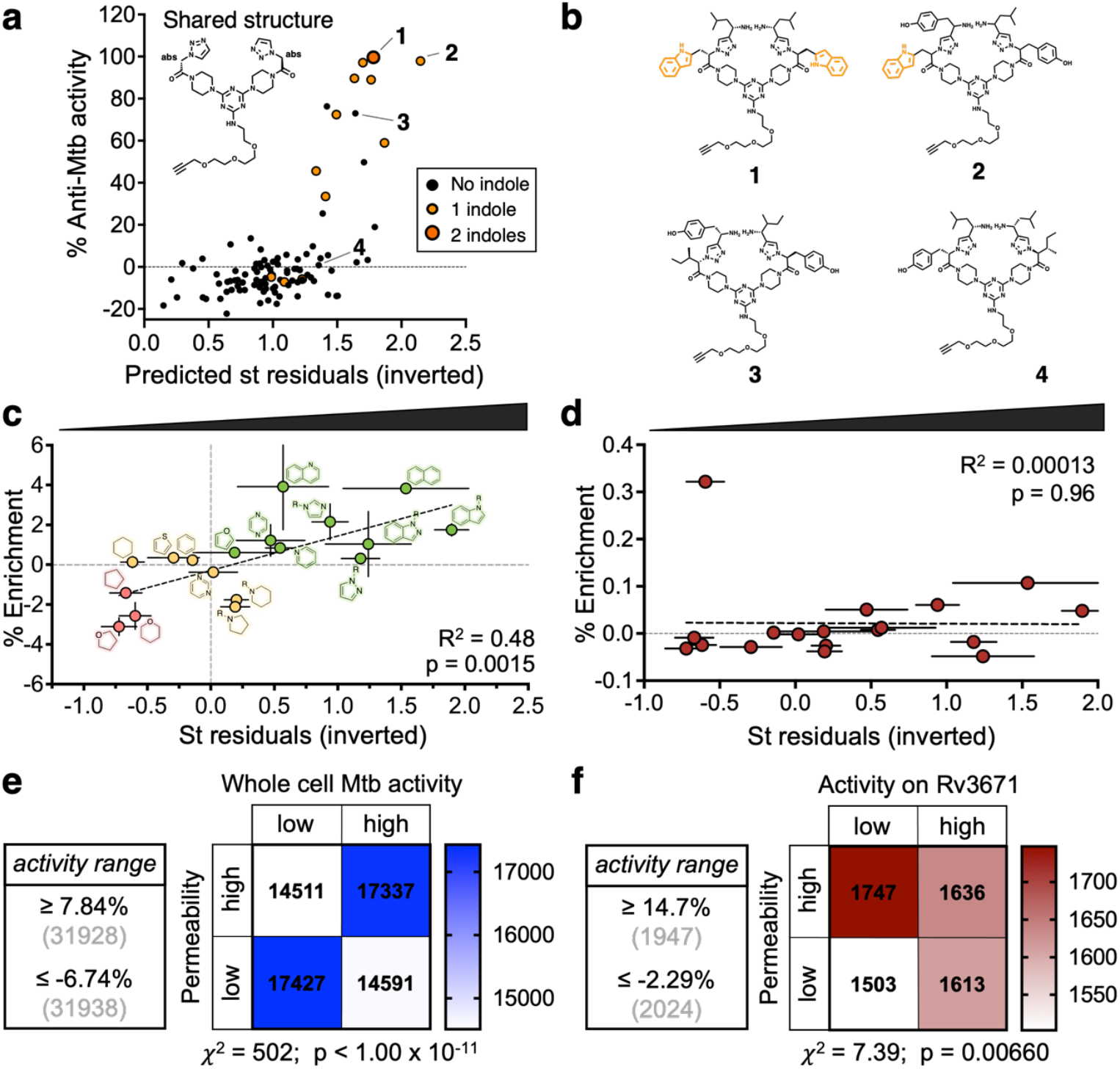
Scaffolds and chemical features that influence mycomembrane permeation correlate with whole cell anti-Mtb activity. (**a**) Relationship between whole cell anti-Mtb activity and ML-predicted mycomembrane permeability for 101 MLSMR compounds that share the indicated structure. The presence of an indole is highlighted in orange. (**b**) Four examples of structures from (**a**), one for each combination +/-indole and +/-activity. (**c-d**) Scaffold-specific relationship between activity against whole Mtb cells (**c**) or against a purified Mtb enzyme (serine protease Rv3671c (**d**)) and observed mycomembrane permeation (inverted standardized residuals). X axis is median mycomembrane permeability for test azides that share indicated scaffolds (from **Fig. 2a**). Y axis is percentage of activity enrichment for MLSMR and TAACF molecules that share the same scaffold relative to all the molecules in the respective datasets (**Figure S12**). Y axis data available from CDD (**c**) or publicly available from PubChem (AID 2606) (**d**). (**e-f**) Relationship between (**e**) whole cell anti-Mtb activity (MLSMR) or (**f**) purified Rv3671c enzyme inhibition and ML-predicted mycomembrane permeation. Each library was divided into quadrants (threshold for permeability: median; threshold for activity: bottom and top 15% of the total values of the libraries (MLSMR) or bottom and top 1% values (Rv3671c), see **Figure S12** for further analyses) and the number of values per quadrant were plotted in (**e**) and (**f**) respectively. The association in (**e**) is significantly stronger than the association in (**f**) by Cramer’s V.

Because we do not know the targets of these compounds, we next examined the relationship between chemical structure, mycomembrane permeability, and whole cell activity more broadly. We widened our analysis to three different screens: the MLSMR (above) and Tuberculosis Antimicrobial Acquisition and Coordinating Facility (TAACF; data obtained from CDD^35^) small molecule collections, which were screened against whole Mtb cells, and a third screen against the purified Mtb enzyme Rv3671c (PubChem AID 2606). In addition to activity data, the libraries afforded us the opportunity to examine molecules that do not have azide tags and occupy broader chemical space than that covered by our screening libraries (**Fig. S11**). We found that molecule activity in the whole cell screens, but not the enzyme screen, correlates with both observed permeability of different chemical scaffolds (**Fig. 5c-d**; **Fig. S12**) and with ML-predicted mycomembrane permeability (**Fig. 5e-f**; **Fig. S12**). These data suggest that scaffolds and other chemical features that influence mycomembrane permeation are correlates of whole cell anti-Mtb activity.

## Discussion

Mass spectrometry studies that assess whole cell association have found that molecule accumulation varies across *E. coli, P. aeruginosa*, and *Acinetobacter baumannii*, likely reflecting species-specific differences in molecule transit via porins, diffusion across the outer membrane, and transit via efflux pumps^9, 10, 40^. Given that the Mtb mycomembrane has a structure^6, 12, 13^ and porin-like proteins^41, 42^ that are distinct from those of the Gram-negative outer membrane, it is not surprising that the predictors of mycomembrane permeation that we identify here are different from those of Gram-negative bacteria accumulation^9, 10, 27^. Future use of Mtb strains with targeted disruptions to mycomembrane transporters or structural integrity may enable the identification of pathway-specific predictors of permeation.

One limitation of PAC-MAN is that it requires the presence of an azide. Azides are widely recognized as ideal bioorthogonal tags because they are small and have minimal known impacts on the physicochemical properties of the parent compound^21, 43^. While we cannot rule out potential effects of the azide on mycomembrane permeation, we note that the chemical features that we identified as predictors of this phenotype from azide-tagged libraries are also correlates of whole cell activity for untagged compounds (**Fig. 5**; **Fig. S10**; **Fig. S12**). We speculate that the constancy of the azide across all the test molecules, the large number of test molecules, and the redundancy of some test molecules (*i*.*e*., compounds that differ only in the position of the azide) contribute to our ability to predict mycomembrane permeation.

Intracellular accumulation in Mtb depends on the ability of a molecule to overcome membrane, efflux, and metabolism barriers^2, 3^. Because it covalently traps test molecules in the mycobacterial cell wall, PAC-MAN primarily measures mycomembrane permeation, *e*.*g*., passive diffusion and facilitated transport^7, 20^. As well, our test molecules are relatively simple in structure as they have a median size of 180 Da and generally abide by Lipinski’s rule of five^44^. The restrictions on cell compartment and chemical space queried are both a strength and limitation of our work. On the one hand, we were able to identify clear structure-function relationships for an aspect of intracellular accumulation. Moreover, these relationships, where tested, accurately predicted mycomembrane permeation. Chemical predictors of mycomembrane permeation have obvious potential in informing the (re)design of anti-Mtb therapeutics against periplasmic targets. Direct and indirect data from this work and the literature, respectively, suggest these predictors may have even broader utility, *i*.*e*., evaluation of compounds with cytoplasmic targets. On the other hand, the structure-function relationships that we identify here may not suffice as predictors for larger, more complex molecules or for whole cell accumulation.

Expansion of the chemical space covered by our azide libraries beyond Lipinski’s rule of five^45^, along with whole cell mass spectrometry^4, 5, 9^-^11, 40, 46, 47^, adaptation of PAC-MAN for the cytoplasm^48^, and the use of strains with defined membrane, efflux, or metabolism defects^4, 48, 49^ will collectively enable the generation of more comprehensive accumulation models.

## Acknowledgments

We thank the director of the University of Massachusetts Amherst Flow Cytometry facilities, Dr. Amy Burnside, for her help and advice. These studies were supported by the National Institutes of Health (R01 AI179080 to W.I., M.M.P., and M.S.S. as well as U19 AI142731 and R01 AI153145 to JSF), by the University of Massachusetts Amherst Institute for Applied Life Sciences Midigrant and Core Facilities Incentive Funds to I.L., and by the Gates Foundation INV-080847 to M.M.P. and M.S.S.. This work utilized resources from Unity, a collaborative, multi-institutional high-performance computing cluster managed by UMass Amherst Research Computing and Data.

## Author contributions

I.L., Z.L., N.E., S.F. equally contributed to this work; I.L., Z.L., N.E., S.F., J.S.F., W.I., A.G.G., M.M.P., and M.S.S. designed research; I.L., Z.L., N.E., S.F., T.P.B., K.M., S., M.W., A.G., and T.G. performed research; I.L., Z.L., N.E., S.F., T.P.B., K.M., S., M.W., A.G., T.G., J.D., J.S.F., W.I., A.G.G., M.M.P., and M.S.S. analyzed data; J.D., J.S.F., W.I., A.G.G., M.M.P., and M.S.S. supervised the research; I.L. and M.S.S. wrote the manuscript with the help of all authors. The manuscript was discussed and approved by all authors.

## Supplementary Information is available for this paper

## Materials Availability

All reagents generated in this study are available upon request from the corresponding authors.

## Code and Data Availability

All code, processed input data, and saved model weights for the cheminformatics analyses and MycoPermeNet model are available upon request and will be made publicly available prior to final publication on github: https://github.com/Nevbarunegbe/Mycomembrane-permeability-project.

## Material and methods

For a full description of the experimental procedures see **Supplemental Methods** file.

