## Supplementary Information for "Identification of chemical features that influence mycomembrane permeation and antitubercular activity"

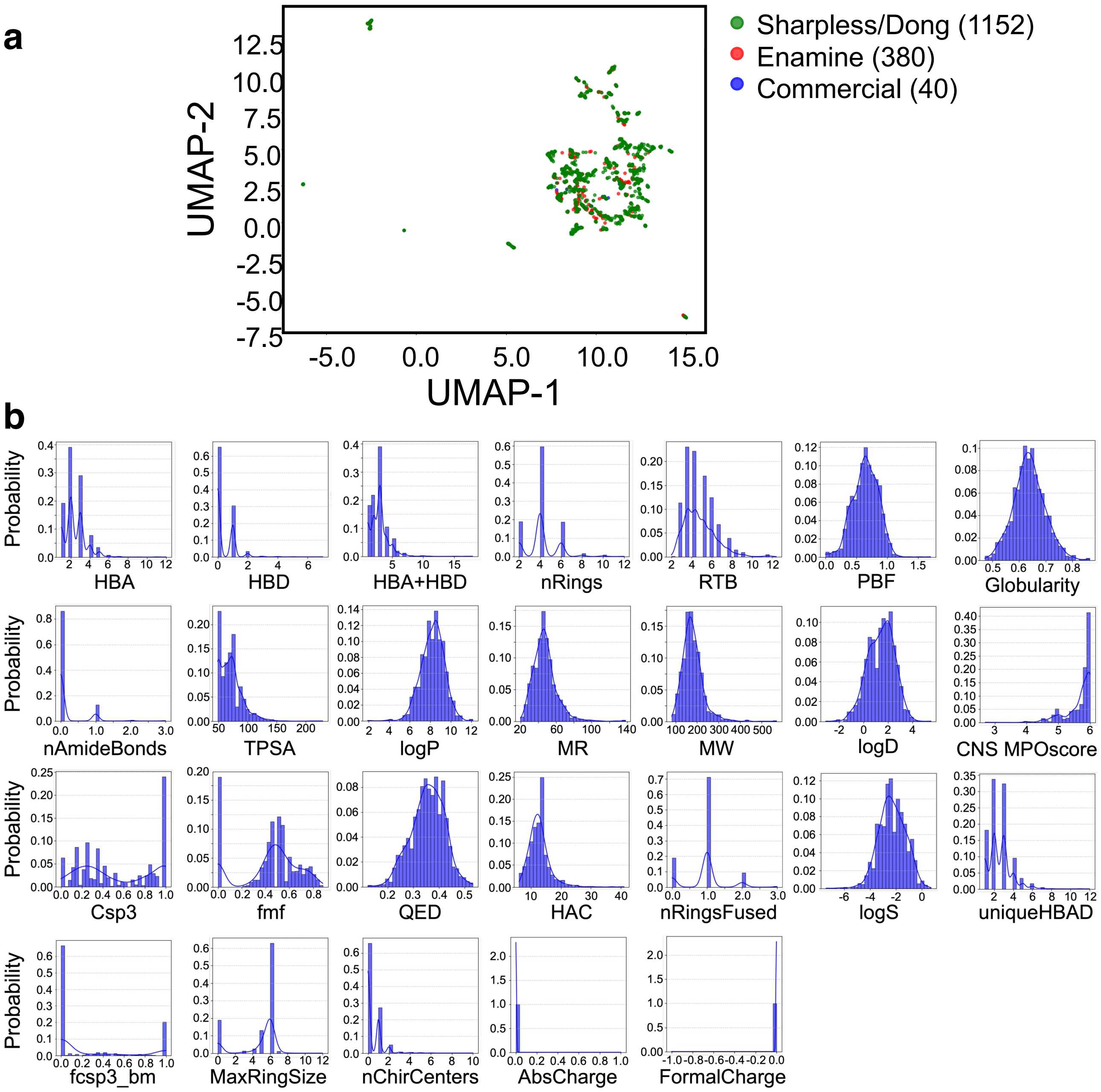


**Figure S1. Chemical features of azide-tagged molecules screened by PAC-MAN.** (**a**) 2D graph representing the chemical space of the three libraries (Sharpless/Dong, Enamine, and commercial) was generated by using the Uniform Manifold Approximation and Projection (UMAP) algorithm. (**b**) Distribution of major physicochemical properties of the libraries. HBA: hydrogen bond acceptor; HBD: hydrogen bond donor; nRing: number of rings; RTB: number of rotatable bonds; TPSA: topological polar surface area (Å$^2$); Log P: *n*-Octanol/water partition coefficient (log); MR: Molar Refractivity (Å$^3$); MW: molecular weight (Da); Csp3: fraction of sp$^3$ carbons; fmf: fraction of heavy atoms in the molecular scaffold; QED: Quantitative estimation of drug-likeness; HAC: heavy atom count; nFusedRings: number of fused rings; uniqueHBAD: number of unique hydrogen bond donors and acceptors; MaxRingSize: Maximum Ring Size; fcsp3_bm: fraction of Csp$^3$ hybridized carbons in the molecular scaffold; PBF: Plane of Best Fit ; AbsCharge: absolute charge; FormalCharge: formal charge; Log D: *n*-Octanol/water distribution coefficient at pH 7.4 (log); Log S: water solubility (log); CNS MPO score: central nervous system multiparameter optimization. Physicochemical properties were calculated using RDKit except for LogD, LogS and CNS MPO score that were calculated with Collaborative Drug Discovery (CDD).

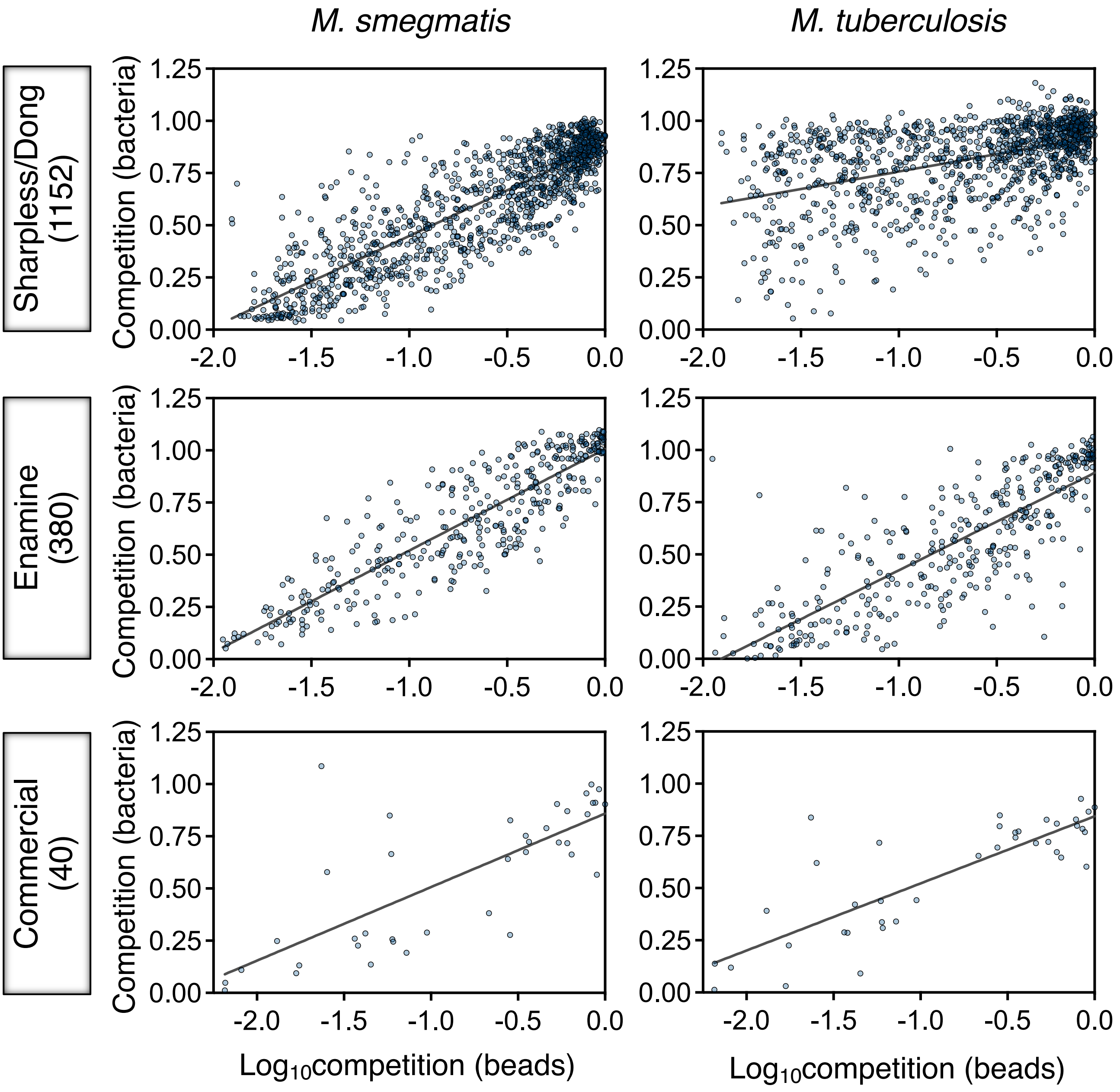


**Figure S2. Log-linear correlation between test azide competition on DBCO-beads vs. DBCO-labeled bacteria.** Screenings on *M. smegmatis* (*left*) and *M. tuberculosis* (*right*) were performed using test azides from Sharpless/Dong (1152), Enamine (380), and other commercial sources (40). Standardized residuals were generated for each library separately to account for potential differences in library quality.

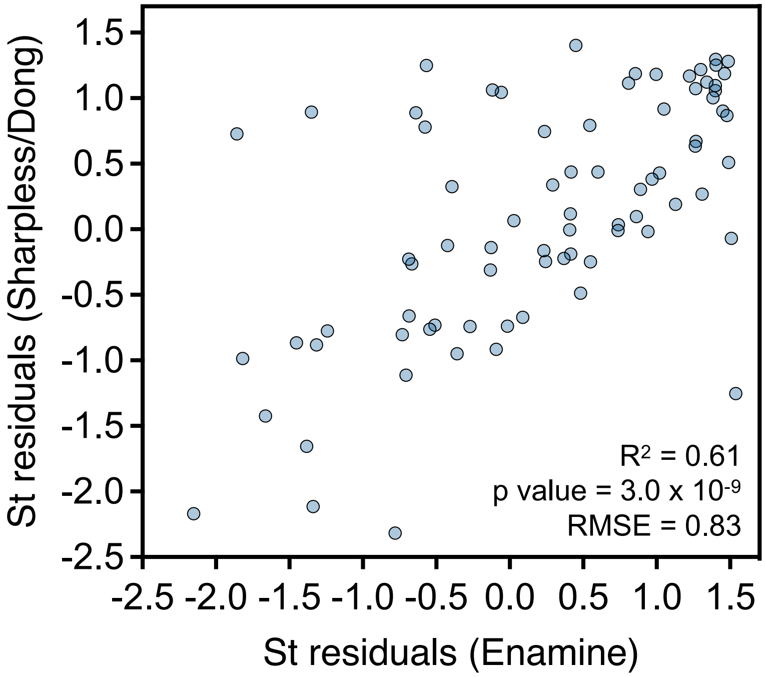


**Figure S3. Consistency of standardized residuals among azide libraries.** Comparison of standardized residuals for test azides that are shared between the Sharpless/Dong (1152) and Enamine (380) libraries. RMSE: Root Mean Square Error.

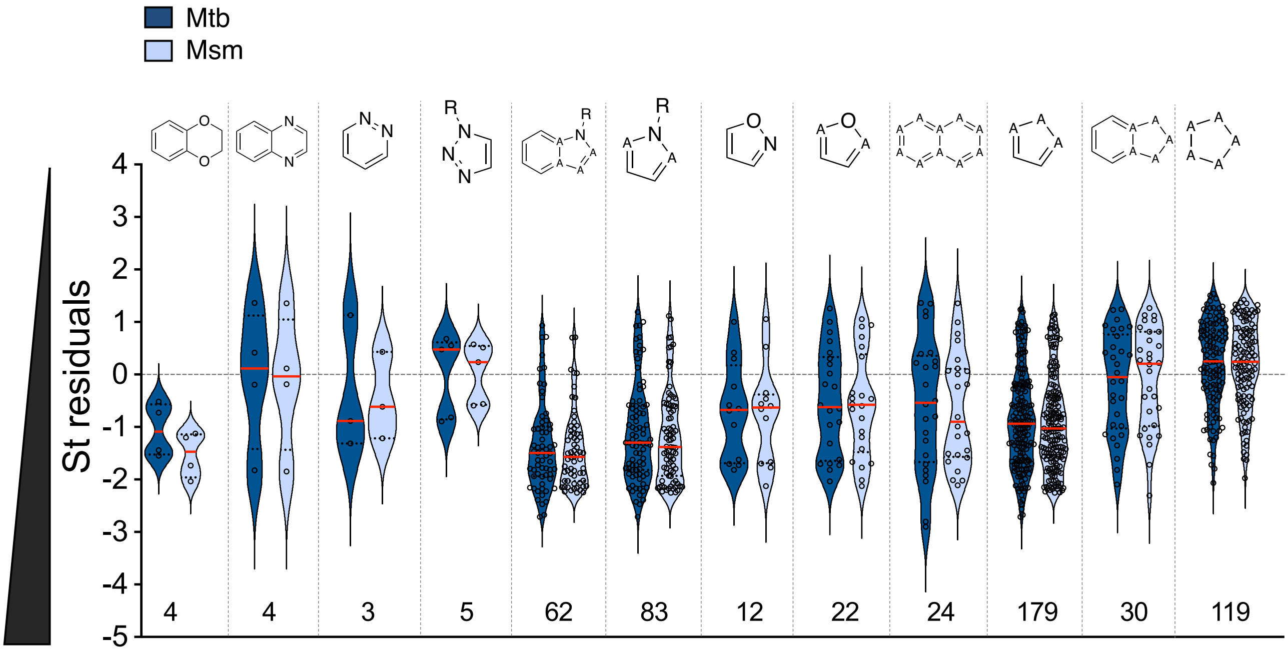


**Figure S4. Expanded scaffold analysis.** Expanded set of chemical scaffolds analyzed as in **Fig. 2a**. The presence of aromatic heterocycles bearing more than one heteroatom positively correlates with mycomembrane permeability. Loss of aromaticity (last two scaffolds) negatively correlates with mycomembrane permeability. A: any atom.

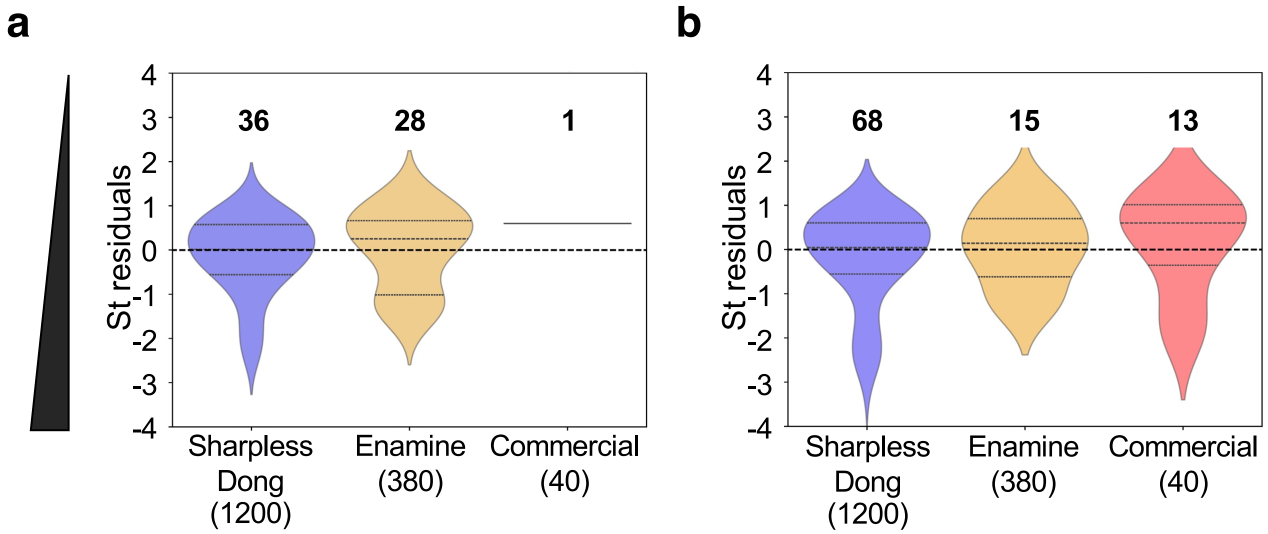


**Figure S5. Chemical properties that correlate with whole cell accumulation in Gram-negative bacteria do not predict mycomembrane permeability.** Mycomembrane permeability scores for test azides that abide by (**a**) *E. coli* eNTRy rules[13] (presence of an ionizable Nitrogen; low Three-dimensionality; low globularity (radius of gyration/molecular volume ≤ 0.25); and Rigid (≤ 5 rotatable bonds)) or (**b**) *P. aeruginosa* PASsagE rules (*Pseudomonas aeruginosa* Self-promoted Entry[12, 27] positive polar surface area (Q_VSA_PPOS ≥ 80); positive formal charge (FC ≥ .98); and high hydrogen bond donor surface area (HBDSA ≥23)). Median values of the standardized residuals are ≥ 0, indicating low permeability in Mtb.

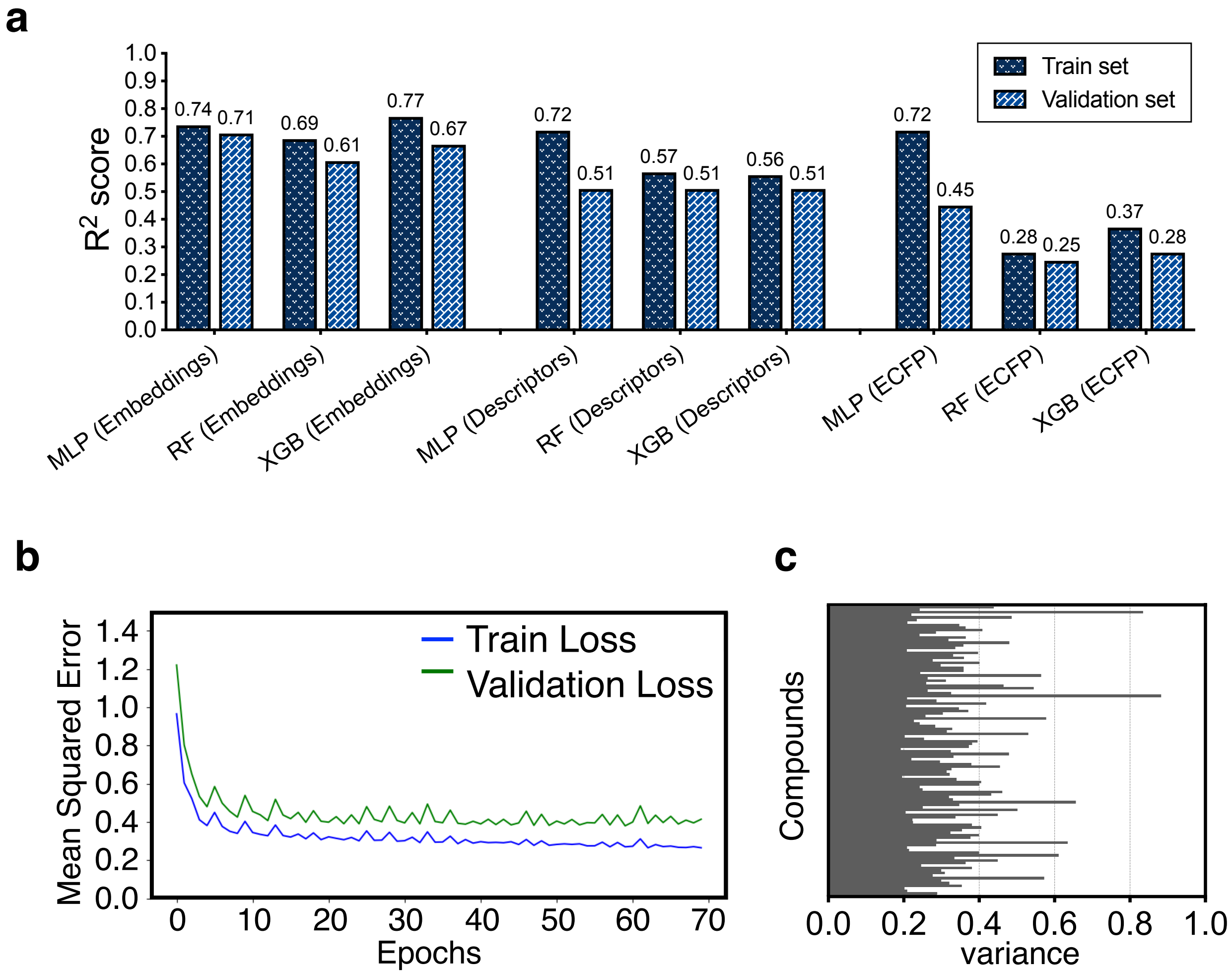


**Figure S6. ML model development and validation.** (**a**) Performance across different ML models. Multilayer perceptron was chosen because it shows the closest performance ($R^2$) in the train and validation sets. (**b**) Trend of ML model for the train and validation set to ensure high performance over epochs and check for data overfitting. (**c**) Uncertainty of the ML model prediction for the test set. ECFP is length 1024 and has bit 3. MLP: Multilayer Perceptron; RF: Random Forest Regressor; XGB: XGBoost Regressor; ECFP: Extended-connectivity fingerprints. See the methods section for more details.

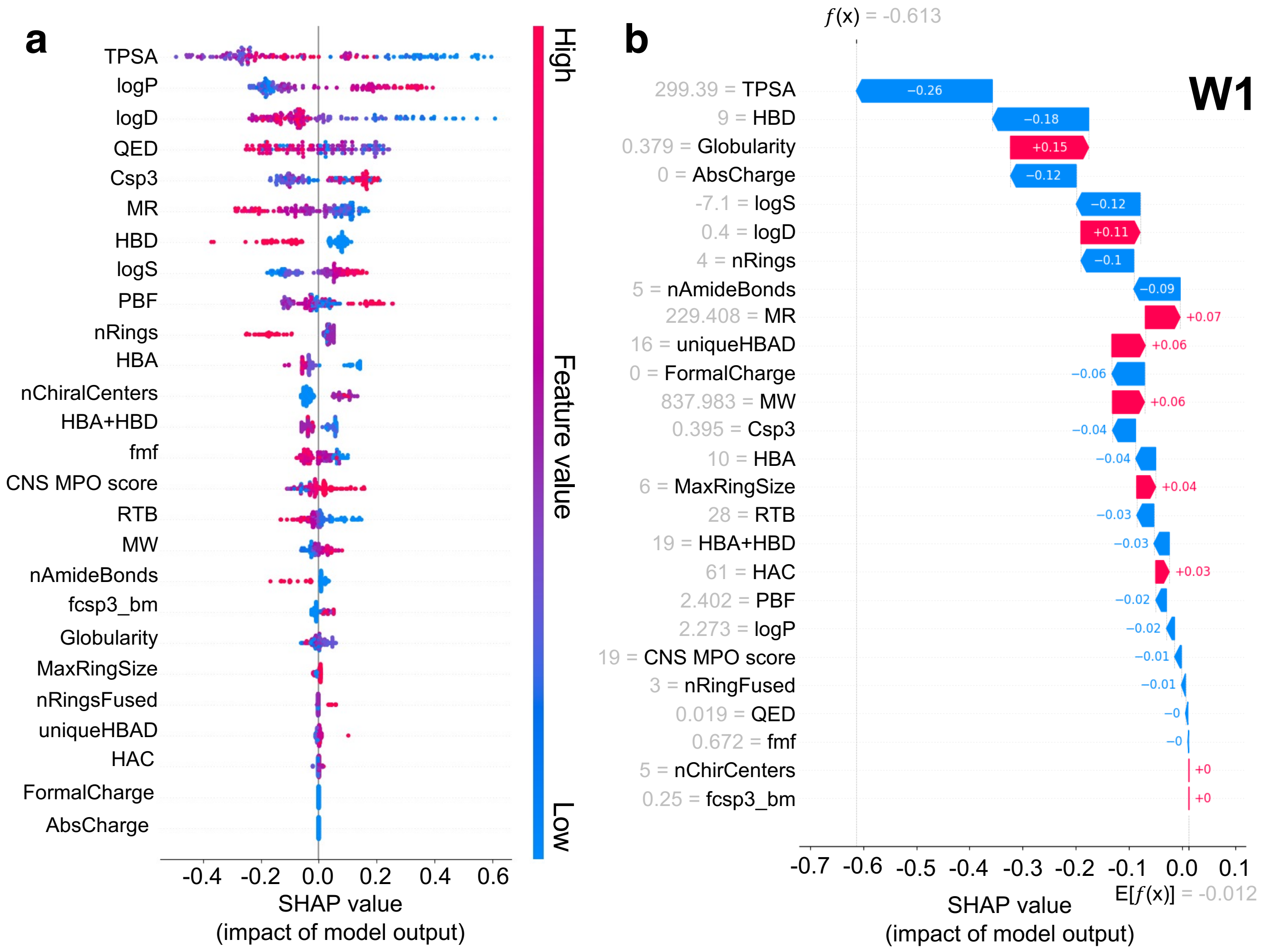


**Figure S7. Interpretability studies of MycoPermeNet** (**a**) Global SHAP (SHapley Additive exPlanations) analysis for the entire azide library; ranks from the top-most impacting feature to the model's prediction from top to bottom. TPSA has the largest impact on the model's prediction. Blue indicates low values of the feature, and red indicates high values of the feature. Points on the left side of the plot show positive impact on Mtb's permeability globally. Each dot represents a compound within the library. (**b**) Local SHAP analysis for an individual compound's prediction (shown here is the compound **W1**, **Fig. 4**). The waterfall plot shows the model contributions of each of the 26 features by pushing the mean of the complex model's predictions (*i.e.,* E[$f$(x)]), either negatively (blue) or positively (red), to the final permeability prediction of that compound (*i.e., $f$*(x)). Blue indicates a positive contribution to high permeability and red indicates a negative contribution. The numbers attached to the features are the exact descriptor values for the **W1** compound.

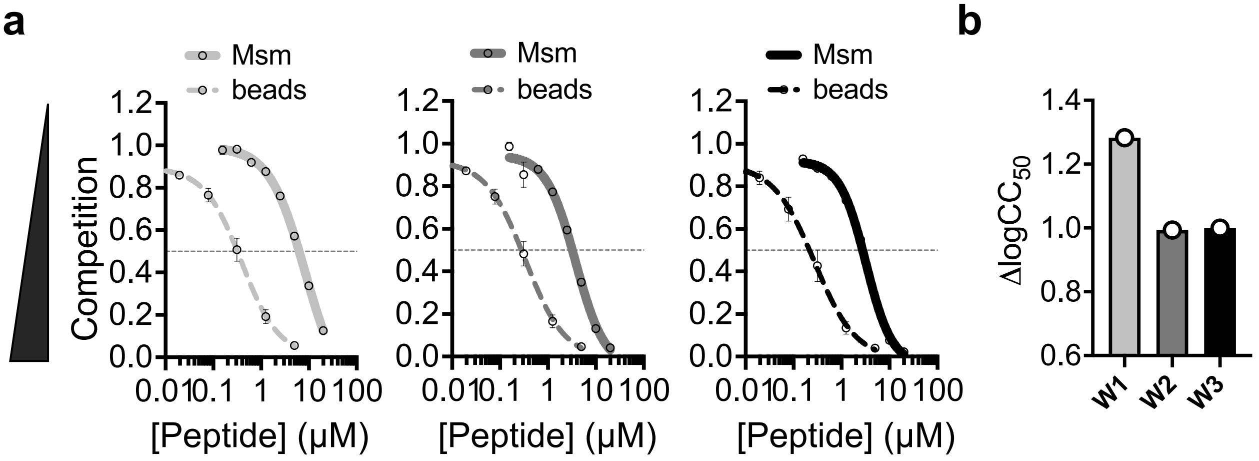


**Figure S8. W1-3 pentapeptide permeability in *M. smegmatis*.** (**a**) PAC-MAN assay for *M. smegmatis* (Msm) and DBCO-bead and (**b**) related $\Delta log_{10}CC_{50}$ as measure of mycomembrane permeability.

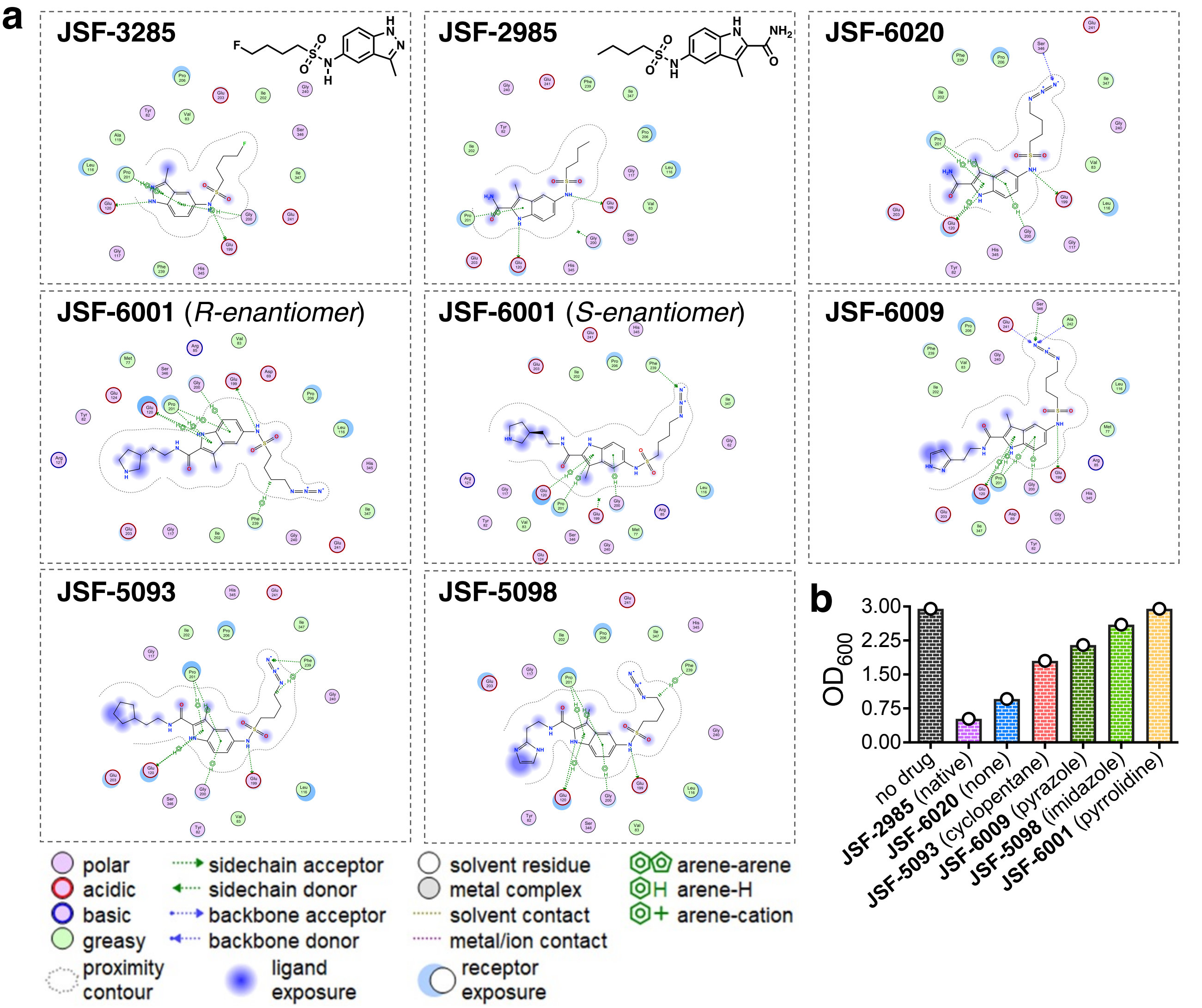


**c**

| Compound | Side Chain | MIC (μM) Mtb H37Rv | RMSD(Å) with respect to JSF-3285 crystal structure | H-bond with Glu199 | H-bond with Glu120 | Arene-H interaction with Pro201 |
|---|---|---|---|---|---|---|
| **JSF-2985** | 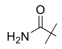 | 3.1 | 1.0 | Yes | Yes | Yes |
| **az-JSF-2985 (JSF-6020)** | 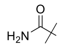 | 25 | 0.94 | Yes | Yes | Yes |
| **az-JSF-2985-imidazole (JSF-5098)** | 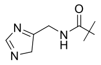 | 12.5 | 2.4 | Yes | Yes | Yes |
| **az-JSF-2985-pyrazole (JSF-6009)** | 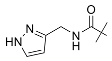 | N/A | 1.8 | Yes | Yes | Yes |
| **az-JSF-2985-pyrrolidine (JSF-6001** ***S-enantiomer*****)** | 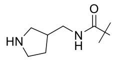 | 100 (*racemate*) | 1.9 | No | Yes | Yes |
| **az-JSF-2985-pyrrolidine (JSF-6001** ***R-enantiomer*****)** | 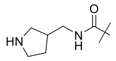 | 100 (*racemate*) | 1.4 | Yes | Yes | Yes |
| **az-JSF-2985-cyclopentane (JSF-5093)** | 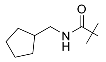 | ≥ 25 (precipitation at 50 μM) | 2.8 | Yes | No | Yes |

**Figure S9. Docking studies of JSF-2985 series with KasA.** See next page for full legend.

**Figure S9. Docking studies of JSF-2985 series with KasA.** (**a**) To evaluate potential target interactions of **JSF-2985** azide analogs, molecular docking studies were performed using the X-ray crystal structure of the mycobacterial beta-ketoacyl-ACP synthase I (KasA) with bound inhibitor **JSF-3285** (PDB ID: 6P9L). Informed by this X-ray crystal structure, we prioritized three interactions in evaluating the docked poses of prepared azide compounds that are derivative of **JSF-2985** – a **JSF-3285** analog which has been shown to inhibit *M. tuberculosis* growth (MIC = 3.1 μM). The first two proposed interactions involve hydrogen bonding of the sulfonamide N-H and the indazole/indole N-H with Glu199 and Glu120, respectively. The third proposed interaction is an arene-H interaction between the indazole/indole ring and Pro201. To further validate the docking poses of the analogs, they were overlaid with **JSF-3285** and the root mean square deviation (RMSD) was calculated. Binding poses that deviated greater than 2.5 Å from the **JSF-3285** X-crystal structure pose and did not participate in the three interactions mentioned above were predicted to be less likely to bind the protein. Such studies suggest that all the investigated **JSF-2985** derivatives have a high probability of retaining the ability to bind their target, although the analysis doesn't allow to provide details of the strength of target binding, and thus, activity on the enzyme. (**b**) Mtb mc$^2$6206 whole cell activity of **JSF-2985** derivatives at 30 μM expressed as optical density ($OD_{600}$) after 6 days of treatment. (**c**) Summary of docking studies features and activity against Mtb H37Rv of **JSF-2985** derivatives. MIC: minimum inhibitory concentration; ND: not determined.

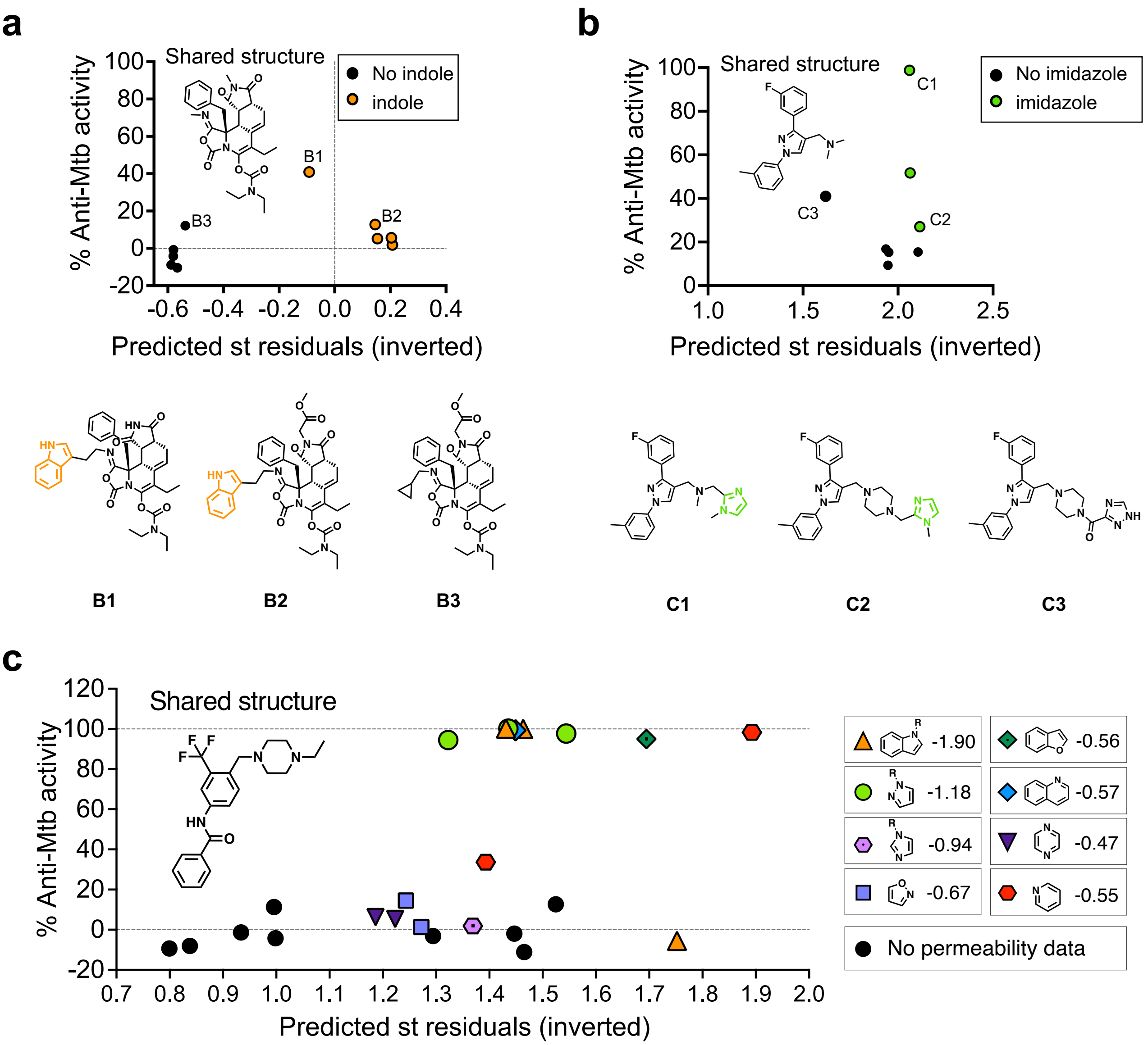


**Figure S10. Scaffolds that influence mycomembrane permeation correlate with whole cell anti-Mtb activity.** (**a-c**) Three additional sets of compounds subjected to the same analysis as in **Fig. 5a**. (**a-b**) The presence of indole (orange, (**a**)), or imidazole (green, (**b**)) which are associated with higher mycomembrane permeability (**Fig. 2a-b**), correlates with higher ML-predicted mycomembrane permeability and whole cell activity. (**c**) The presence of scaffolds (various colors) that are associated with higher mycomembrane permeability (**Fig. 2a-b**) correlates with higher ML-predicted mycomembrane permeability and whole cell activity. The median mycomembrane permeability for each scaffold (standardized residuals) is shown to the right; lower values indicate higher mycomembrane permeability.

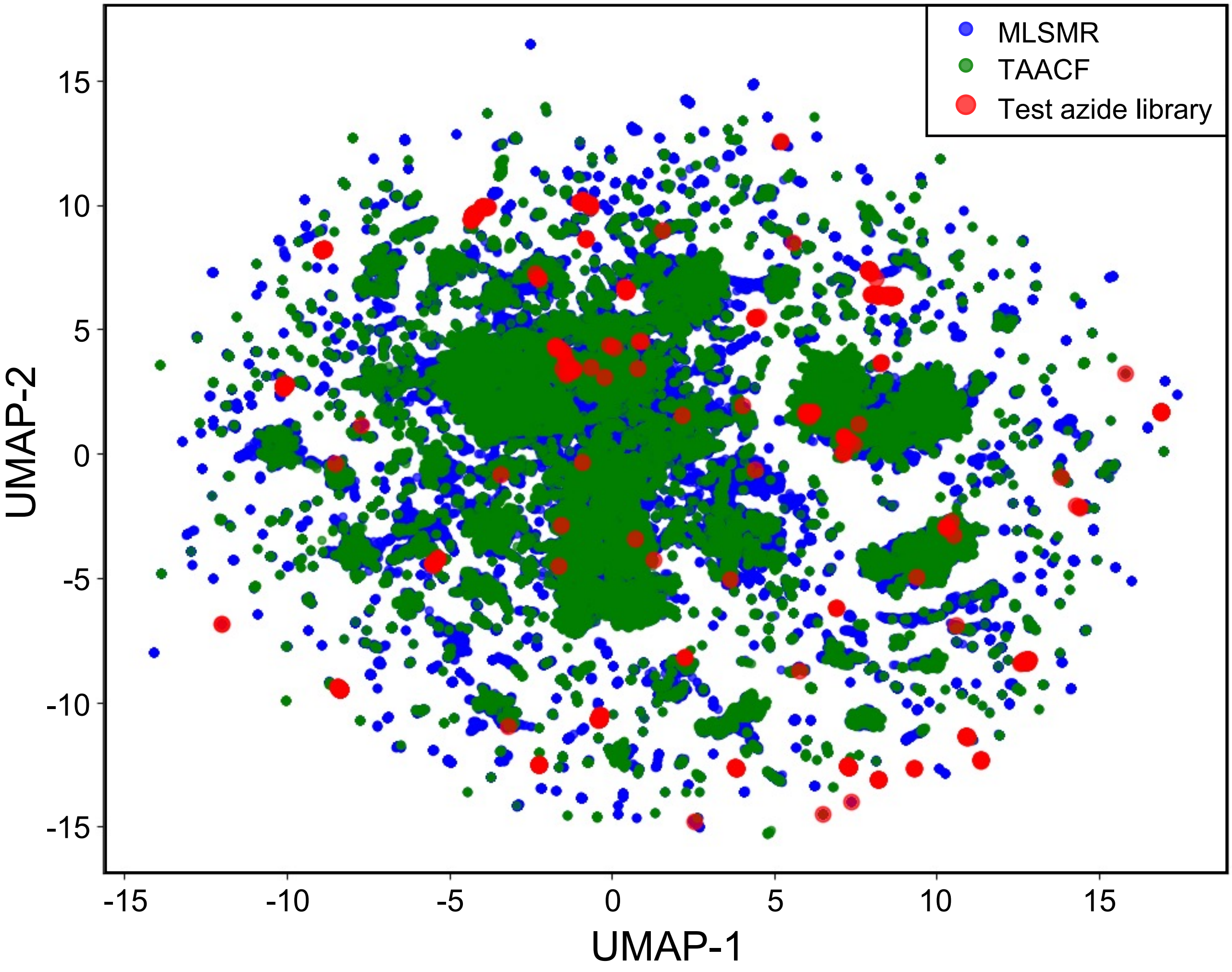


**Figure S11. Chemical space of MLSMR and TAACF collections compared to test azide libraries.** 2D graph representing the chemical space of the libraries was generated by using Uniform Manifold Approximation and Projection (UMAP) algorithm.

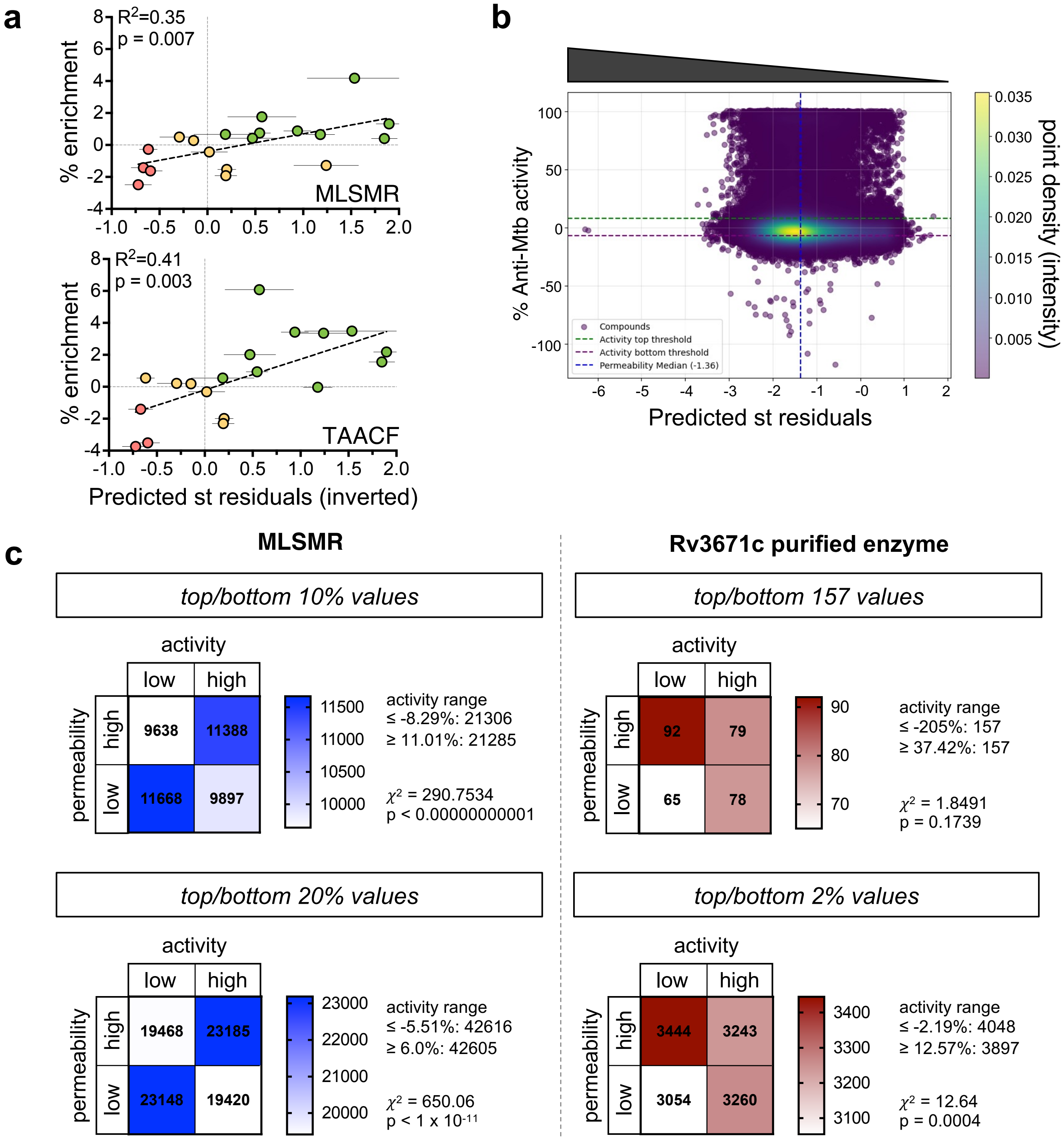


**Figure S12. Correlation between predicted mycomembrane permeation and whole cell anti-Mtb activity.** (**a**) Scaffold-specific relationship between whole cell activity and observed mycomembrane permeation (inverted standardized residuals) shown separately for MLSMR and TAACF screens (merged in **Fig. 5c**). (**b**) Graphic representation of the analysis performed on MLSMR in **Fig. 5e** and **Fig. S12c**. MLSMR library data were graphed as ML-predicted permeability (standardized residuals) vs. anti-Mtb activity then divided in quadrants. Threshold for permeability: median. Threshold for activity: 15% top value and 15% bottom values. (**c**) Extended analysis of **Fig. 5e-f** with additional thresholds as specified in the panels. For the screening on Rc3671c the threshold of activity 37.42% (157 compounds) was defined by the screening's guidelines (PubChem AID 2606).
