## Supplemental Methods for "Identification of chemical features that influence mycomembrane permeation and antitubercular activity"

### Material and methods

#### 1. Experimental procedures

#### Bacterial strains and growth conditions

The double auxotroph *M. tuberculosis* (*Mtb*) mc$^2$6206 strain (H37Rv Δ*panCD* Δ*leuCD*[1]) was provided by Dr. William Jacobs and was cultured in Middlebrook 7H9 supplemented with 0.5% glycerol, 10% Middlebrook Oleic Albumin Dextrose Catalase (OADC; BD), and either 0.05% Tyloxapol, for strain maintenance, or 0.05% Tween-80, for experimental manipulation. Growth media were additionally supplemented with 50 µg/ml L-leucine and 50 µg/ml pantothenic acid. Bacterial stocks were prepared by directly freezing at -80°C *Mtb* culture at OD600 ~0.5.

*Mycobacterium smegmatis* mc$^2$155 [2, 3] were provided by Dr. Anil Ojha and cultured in Middlebrook 7H9 supplemented with 0.5% glycerol (Fisher), 0.05% Tween-80 (Sigma), 10% ADC (ADC: 5% bovine serum albumin, 2% dextrose, 0.003% catalase in water; alternatively catalase is not included but NaCl 0.9% is added). All bacteria were grown 37$^o$ C shaking.

#### High throughput permeability assay (PAC-MAN) on live *in vitro* cultured *M. tuberculosis* mc$^2$6206

*M. tuberculosis* was diluted to $OD_{600}$ ~0.015 and grown in 7H9 +/- 25 μM **TetD** (WuXi AppTec) for 4 days until $OD_{600}$ ~0.25, washed twice with the same medium then incubated with 50 μM of azide-test molecule in 7H9 in 96-well plates for 2 hours at 37 °C shaking. Azide-test molecules were removed by spinning and supernatant removal and bacteria were then resuspended in 7H9 containing 17 μM of the fluorogenic label CalFluor647-azide (APC flow cytometry channel) for 1 hour at 37 °C shaking. The fluorophore was removed by spinning and supernatant removal and bacteria were fixed with fresh prepared 4% paraformaldehyde (PFA, Ted Pella) in polyphosphate buffer (PBS, Genesee Scientific) for at least 2 hours at room temperature. Then bacteria were resuspended in physiological solution (0.9% NaCl in water) and analyzed by BD DUAL LSRFortessa.

#### High throughput permeability assay (PAC-MAN) on live *M. smegmatis*

*M. smegmatis* was inoculated from the glycerol stock by 1 to 1000 dilution to 50 mL fresh 7H9 media with ADC in 250 mL Erlenmeyer flasks each day and grown for 24 hours until 0.5-0.6 OD. 25 μM **TetD** were added to the media, and the cells were grown overnight to achieve stationary phase. The cells were harvested and spun down for 10 min at 3000 g, washed with 50 mL PBST two times and resuspended in 50 mL PBST. Each day, the azido compounds were prepared at 50 mM in PBST from their DMSO stocks. To 4 96-well plates were added 90 μL cells, which were then spun down at 2700 g for 2 min with the supernatant decanted to afford the cell pellets in each well. 100 μL 50 μM of each azido compounds were added to each well with the cell pellet. The plates were incubated in 37 ℃ for 2 h. The cells were spun down for 2 min at 2700 g. The supernatant was decanted, and 100 μL 50 μM Az-6-FAM (Lumiprobe, FITC flow cytometry channel) was then added to each well, followed by incubation in 37 ℃ for 1 h. The cells were then spun down again, and the pellets were washed with 200 μL PBST once and fixed with 100 μL 4% formaldehyde for 15 min. The samples were then analyzed by Attune™ NxT Acoustic Focusing Cytometer.

Alternatively, Msm was grown in 7H9 +/- 25 μM **TetD** for 2 days until $OD_{600}$ ~0.4, washed twice with PBST and incubated with 50 μM of azide-test molecule in PBST in 96-well plates for 2 hours at 37 °C shaking. Azide-test molecules were removed by spinning and supernatant removal and bacteria were then resuspended in PBST containing 1 μM of the fluorogenic label CalFluor647-azide for 1 hour at 37 °C shaking. The fluorophore was removed by spinning and supernatant removal and bacteria were fixed with fresh prepared 2% paraformaldehyde in PBS, then resuspended in physiological solution (0.9% NaCl in water) and analyzed by BD DUAL LSRFortessa. Values normalization: first, fluorescence values of no-**TetD** controls were subtracted from all the samples. The adjusted fluorescence values were then divided by fluorescence values from no-test azide (vehicle, DMSO) control bacteria. Results were concordant between protocols.

**PAC-MAN competition curves of W1-3 peptides and JSF-2985 analogues on live *in vitro* cultured *M. tuberculosis* mc²6206**

*M. tuberculosis* was diluted to $OD_{600}$ ~0.015 and grown in 7H9 +/- 25 μM **TetD** for 4 days until $OD_{600}$ ~0.25, washed twice with 7H9, then incubated with various concentrations of azide-test molecule (or native **JSF-2985**) in 7H9 in 96-well plates for

2 hours at 37 °C shaking. Azide-test molecules were removed by spinning and supernatant removal and bacteria were then resuspended in 7H9 containing 17 μM of the fluorogenic label CalFluor647-azide for 1 hour at 37 °C shaking. The fluorophore was removed by spinning and supernatant removal and bacteria were fixed with fresh prepared 4% PFA for at least 2 hours at room temperature. Then bacteria were resuspended in physiological solution (0.9% NaCl in water) and analyzed by BD DUAL LSRFortessa. Fluorescence values for W1-3 peptides were normalized as above. Given the permeabilization property of **JSF-2985**, the fluorescence for **JSF-2985** analogues was normalized over the bacteria treated with the native (azide-free) **JSF-2985** compound (instead of just the vehicle) for each concentration.

**PAC-MAN competition curves of W1-3 peptides and small molecules on live *M. smegmatis***

DBCO-modified *M. smegmatis* was generated as above. Compounds in different concentrations were prepared on separate 96-well plates. To each well of a 96-well plate was added 90 μL of DBCO-modified *M. smegmatis* followed by spinning down for 2 min at 2700 g, after which the supernatant from each well was decanted to afford the cell pellets. 100 mL compounds in desired concentrations were added to the cell pellets in the working 96-well plate. Each test azide was assayed in technical triplicate. The plates were incubated in 37 ℃ for 2 h. Bacteria were spun down for 2 min at 2700 g. The supernatant was decanted, and 100 μL 50 μM Az-6-FAM was then added to each well, followed by incubation in 37 ℃ for 1 h. Bacteria were then spun down again, and the pellets were washed once with 200 μL PBST then fixed with 100 μL 4% formaldehyde for 15 min. Samples were analyzed by Attune™ NxT Acoustic Focusing Cytometer.

**DBCO modification of polystyrene beads**

1 mL amino functionalized polystyrene beads (5% w/v, 5 mg) were spun down at 21000 g for 15 min in a 1.7 mL microcentrifuge tube and washed with 1 mL deionized water before use. The beads were then spun down and resuspended in 10 mL 100 mM sodium borate buffer pH 9 with 1 μg/mL DBCO-NHS and reacted in 37 °C for 2 h with shaking. The resulting beads were spun down at 4000 g for 10 min, washed once and resuspended in sodium borate buffer. 200 μL acetic anhydride was added to the

suspension and reacted in 37 °C for 2h with shaking. The resulting product was then spun down and washed twice with 10 mL sodium borate buffer and twice with 10 mL PBS and resuspend in 20 mL PBS for further use.

**Competition with the test molecules on DBCO-modified polystyrene beads**

DBCO-modified polystyrene beads were 1 to 20 diluted in PBS before adding to the assay. To a MultiScreen HTS 96-well filter plate with 0.65 mm hydrophilic PVDF membrane added 100 μL beads each well, which was then filtered under vacuum to get the pellet of beads on the filter membrane. 50 μM test molecules in the library were added to the bead's pellets in each well with a final volume of 100 μL, which were resuspended with volumetric pipettes. The plates were incubated in 37 °C for 2 h and then washed with 200 mL PBS, filtered under vacuum to remove the supernatant. 100 μL 50 μM Az-6-FAM was then added to each well. The plates were then incubated in 37 °C for 1h. The beads were washed with 200 μL PBS twice by vacuum filtration and then resuspended in 200 μL PBS. The samples were then analyzed by Attune™ NxT Acoustic Focusing Cytometer.

Alternatively, DBCO-modified polystyrene beads were 1 to 20 diluted in PBST and incubated with the proper concentration of azide-test molecule in 96-well plates for 2 hours at 37 °C shaking. Azide-test molecules were removed by spinning and supernatant removal and beads were resuspended in PBST containing 1 μM of the fluorogenic label CalFluor647-azide for 1 hour at 37 °C shaking. The fluorophore was washed and resuspended in physiological solution and analyzed by BD DUAL LSRFortessa. Results were concordant between protocols.

**JSF-2985 analogues growth inhibition on *M. tuberculosis* mc$^2$6206**

*M. tuberculosis* was diluted to $OD_{600}$ 0.01 and incubated with **JSF-2985** analogues (or just vehicle, DMSO) 30 μM for 6 days and $OD_{600}$ was read. 30 μM concentration was chosen because of **JSF-5093** precipitation at higher concentrations.

**JSF-2985 analogues minimum inhibitory concentration (MIC) on *M. tuberculosis* H37Rv**

Determination of MIC values was done using microplate Alamar blue assay (MABA). Serial dilutions of test compounds in 100 μL were prepared in growth media (7H9-ADS

– albumin-dextrose-sodium chloride) in a 96-well microplate (Fisher #FB012932). Each well was added with 100 µL of *M. tuberculosis* culture (H37Rv strain diluted 1:1000 from an initial $OD_{595}$ of 0.2). After 7 days incubation at 37°C, alamarBlue HS Cell Viability Reagent (Invitrogen #A50101) supplemented with 20% Tween 80 was added. Plates were incubated for an additional 24 hours followed by absorbance readings at 570 nm (normalized to 600 nm) to assess bacterial viability[4].

### 2. In silico analysis and experiments

#### *2.1 Cheminformatics*

**Data Cleaning**

To reduce the effect of DBCO bead reactivity on the apparent accumulation, the residual of apparent accumulation against the logarithm of bead reactivity for each molecule was calculated. First, a simple linear regression was performed using NumPy.polyfit() with a degree of one. Then, residuals were calculated according to Equation 1:

Eq. 1

$$r = y_i - \hat{y}_i$$

In Equation 1, $y_i$ refers to the measured accumulation, and $\hat{y}_i$ refers to the accumulation predicted by the line-of-best fit from the linear regression. Next, residuals were standardized according to Equation 2 in order to compare the residuals of different compound libraries.

Eq. 2

$$r_{std} = (r - \mu)/\sigma$$

In Equation 2, the standardized residual for a compound, $r_{std}$ is equal to the original residual, $r$, minus the mean residual among all compounds of that compounds library, $\mu$ divided by the standard deviation among all compounds of that compounds library, $\sigma$. This equation is also known as the z-score of the residual. To combine libraries, we saved the standard residuals in a Pandas data frame, and, looping through each library, added the molecules to a new data frame. If we came across a molecule common among two or more libraries, we took the standardized residual value from the 40 compound or 380 compound library.

**Scaffold analysis**

To analyze how the presence of various ring structures impacts the apparent accumulation, we binned compounds into groups based on whether they had at least one instance of each ring structure in their structure. The ring structures chosen are shown in **Fig. 2a** and **Fig. S5** for which the analysis was performed using Collaborative Drug Discovery (CDD)[5]. For **Fig. 2b** the analysis, to further classify compounds, a distinction was made between ring structures which were part of a larger ring structure (FUSED), and those which were not (NONFUSED). These groupings were then used to plot the distribution of standardized residuals in violin plots with Graphpad Prism.

**Physicochemical Property Correlation analysis**

To analyze how physicochemical properties of the compounds may affect the apparent accumulation, various properties were calculated with the rdMolDescriptors package within RDKit. The properties calculated were Number of Hydrogen Bond Acceptors (HBA), Number of Hydrogen Bond Donors (HBD), HBA + HBD, Number of Rings (nRings), Number of Rotatable Bonds (RTB), Number of Amide Bonds (nAmideBonds), Globularity, Plane of Best Fit (PBF), Topological Polar Surface Area (TPSA), Lipophilicity (logP), Molar Refractivity (MR), Molecular Weight (MW), Number of $Csp^3$ hybridized carbons (Csp3), Fraction of heavy atoms in the molecular scaffold (fmf), Quantitative estimation of drug-likeness (QED), Heavy Atom Count (HAC), Number of Fused Rings (nRingsFused), number of unique hydrogen bond donors and acceptors (uniqueHBAD), Maximum Ring Size (MaxRingSize), Number of Chiral Centers (nChiralCenters), fraction of $Csp^3$ hybridized carbons in the molecular scaffold (fcsp3_bm), Formal Charge (FormalCharge), and Absolute Charge (AbsoluteCharge). We further calculated 3 additional properties from the CDD visualization tool, which are *n*-Octanol/water distribution coefficient at log pH 7.4 (logD), water solubility (logS) and Central Nervous System Multiparameter Optimization (CNS MPO) score, resulting in a total of 26 physicochemical properties. If any properties were not able to be calculated for a compound, that compound was discarded. The correlation matrix for the properties along with raw apparent accumulation, logarithm of bead reactivity, original residuals, and standardized residuals was calculated with the Pandas DataFrame.corr() function. Properties without valid correlation values were discarded. The correlation matrix was visualized as a heatmap with Matplotlib. LogD, logS, MPO were calculated using the CDD visualization tool. The other properties were calculated using RDkit function calls.

**Effect of a permeability-promoting scaffold within molecule sets with common minimal structure.**

The analyses were performed using the Collaborative Drug Discovery (CDD) software.

*Indole*

Indole-containing molecules were searched within the *Molecular Libraries Small Molecule Repository* (MLSMR) library screened for anti-TB activity, applying a bottom molecular weight (MW) threshold of 500 Da (around 4-5x the indole MW: 117.15 Da). We performed a structural search for molecules where indole was present as a peripheral substituent instead of the main core. Groups where indole occupied a central position and, thus, couldn't be removed to perform a comparative analysis +/- indole, were not considered. We prioritized the compounds that showed structural similarities (shared core structure) within the indole-containing subgroup. For each chosen molecule/group, the indole first, and subsequently additional peripheral parts of the molecule, were removed from the structure to identify a "minimal structure" which was then searched for again within the entire MLSMR dataset, defining the group for the +/- indole analysis. This was a reiterative process to obtain as many compounds as possible for each group, until the structure became too small to be unique. Given the lack of information about the molecular targets relevant to the library, sets of molecules were chosen (≥ 7 molecules) to inform structure-activity relationships (SAR) and decrease the likelihood that indole was directly involved in the target engagement. For the same reason, we prioritized structures for which the scaffold of interest represented a small portion of the molecular structure (i.e., one-fifth of the total MW or less). Around 20 minimal core structures were analysed and only the ones that met the above mentioned criteria were selected. All the chosen sets of molecules present examples of drug structures missing the scaffold of interested but showing activity, suggesting the role of the scaffold as auxophore instead of pharmacophore.

*Imidazole*

The same analysis described above for indole was performed for imidazole-containing compounds but the bottom threshold of MW was set as 300 Da, proportionally with imidazole MW 68.8 Da. The results obtained in **Fig. S10c** were obtained using the threshold for imidazole.

### Correlation between predicted permeability and activity on big published datasets

To investigate relationships between permeability and biological activity, we used previously published drug libraries screened for whole cell anti-TB activity (MLSMR[6]), where mycomembrane permeability can affect activity, and the screening against the purified enzyme Rv3671 (PubChem AID 2606) where mycomembrane permeability is not a factor in activity. This analysis aims at testing the hypothesis that less permeable compounds are also less active, and vice versa. Since we do not know the level of correlation between mycomembrane permeability and activity, we were looking for major statistical differences among the datasets where mycomembrane permeability is a relevant factor for activity (MLSMR) vs the one for which is not (purified Rv3671enzyme).

First, we predicted the permeability of the entire MLSMR and Rv3671 libraries using MycoPermeNet. We divided the libraries in quadrants for high vs. low predicted permeability and high vs. low activity. For predicted permeability, we categorized based on the median permeability for each of dataset individually: values below the median were classified as 'high permeability' and those above the median as 'low permeability', due to the inverted nature of our permeability residuals (i.e. -3 is highest permeability and 3 is lowest permeability).

The activity thresholds were defined differently for the two libraries given the nature of the screening and the datasets. Screened libraries activity data distribution:

MLSMR screening (percentage of growth inhibition at 10 μM): mean: 3.525; standard deviation: 18.485; minimum: -117.650; maximum: 105.380; median: -0.130;

Purified Rv3671 enzyme screening (percentage of inhibition at 5.96 μM): mean -2.353; standard deviation: 13.199 minimum: -776.820; maximum: 511.750; median: -0.850

MLSMR anti TB-screening was performed using AlmarBlue reagent[4] while for the in vitro enzyme screening a competitive activity-based protein profiling assay based on fluorescence polarization was used.[7]
Both datasets show a large number of negative values for activity, which is likely due to experimental noise. To obtain a reasonable threshold without generating a strong imbalance between the amount of datapoints for the active and the inactive compounds we chose a defined percentage of the total values, rather than selecting a percentage cutoff. Since the activity ranking among the negative values is meaningless, we chose percentage of values that provided balanced thresholds for high and low active close to the 0.

Both libraries' values were ranked by activity, and the same number of high and low activity values (percentage respect to the whole dataset) were selected: 10%, 15%, and 20% for MLSMR; 157, 1%, and 2% for the purified Rv3671 enzyme screening. The threshold that led to 157 compounds was chosen following the screening directions for the activity threshold (>34.02%).

We carried out a series of statistical tests on both datasets in order to ensure statistical significance. Statistical experiments were done using the Scipy package (version 1.13.1).

After classification into groups, we assessed the association between permeability and biological activity using a contingency table which compares the permeability category (rows) and the activity category (columns). A Chi-Squared Test for independence was performed on this table to determine whether these categories were significantly associated, yielding a chi-squared statistic ($\chi^2$), p-value and degrees of freedom for an all-round assessment (see **Fig. 5e-f** and **Fig. S12c** for details).
While for MLSMR the groups "high permeability/high activity" and "low permeability/low activity" showed a strong enrichment compared to the other two, for the purified Rv3671 enzyme a small enrichment was observed for the high permeability/low activity group.

To further validate association strength to distinguish between the two libraries, we carried out Cramér's V test to evaluate statistical association strengths of the libraries.

For MLSMR library we observed a very strong association relative to the purified enzyme library.

#### Docking studies of JSF-2985 analogues with KasA

The X-ray crystal structure of JSF-3285 bound to the KasA protein was retrieved from the RCSB Protein Data Bank (PDB: 6P9L[8]). The protein structure was prepared for docking simulations using the QuickPrep function in the Molecular Operating Environment (MOE) software package[9]. Structure Preparation was used to correct topological errors in protein residues. Hydrogen atoms were added using Protonate3D and water molecules greater than 4.5 Å away from ligand atoms were deleted. Tethers were installed on binding pocket, ligand, and solvent atoms present in the binding pocket to limit their deviation from the experimentally determined coordinates during simulations. Atoms further away from the binding pocket were fixed to increase efficiency during simulation. Refinement and energy minimization of the protein were performed using the Amber:EHT force field. Ligands were minimized using the MMFF94x force field for small molecules, while keeping the protein fixed. Analogs were built using the Builder function in MOE. For the docking calculations, 30 poses were generated for each ligand and docked using the Triangle Matcher placement function. A combination of the GBVI/WSA ΔG and the London ΔG scoring functions were used. The top five scoring conformations were evaluated by overlaying them with the lowest energy docked JSF-3285 conformation.

### *2.2 Machine Learning*

#### Dataset Preparation

The 40, 380, and 1152 azide libraries were standardized and merged using the standardized residual calculation described above. The compounds were then represented as Simplified Molecular Input Line Entry System (SMILES) strings[10]—a standard textual representation of the compounds used for cheminformatics. Defective SMILES that could not be properly parsed by the RDkit tool[11] were removed, resulting in a final dataset of 1558 samples with corresponding permeability residuals labeled as “Mtb Standardlized Residuals.” Values range from -3 (most permeable) to 3 (least permeable). The dataset was then split into training, validation and test sets using the Bemis-Murcko scaffold-balanced strategy implemented by Chemprop[12, 13]. This procedure ensures that compounds with the same molecular scaffold are not shared across data splits, which can result in overestimating model performance. This resulted in 217 unique scaffolds, with the training, validation, and test sets containing 1246, 155, and 157 samples respectively.

**Embedding Space Generation**

We employed a message passing neural network (MPNN) architecture implemented in the Chemprop framework (version 1.6.1), to generate molecular embeddings. MPNN operates in two phases, the first being the message-passing phase which propagates information across the molecular graphs, ultimately building a neural representation of the whole graph. The other phase is the readout phase in which the neural representation generated from the message passing phase is used to predict the property of interest, in our case, the permeability residuals. This readout phase ensures that the generated embedding space is useful for predicting downstream properties. Chemprop implements a directed message passing neural network, a type of MPNN, which centers the message passing on bonds and gathers information of neighboring bonds[12].

Hyperparameter tuning was carried out after testing various architectures, and the following hyperparameters gave the best performance on the validation set. We selected an MPNN with a depth of 3, where each layer consisted of a series of graph convolutions followed by a ReLU activation function. The MPNN encoder was configured with a hidden size of 300, utilizing a linear layer without bias to map input features (147-dimensional) to hidden representations. The output of the MPNN encoder was further processed through a readout function, consisting of a series of linear layers with ReLU activations, to produce the final molecular embedding.

The model was trained for 30 epochs with a batch size of 50. The initial learning rate was set to 0.0001, and a warm-up period of 2 epochs was applied before transitioning to a maximum learning rate of 0.001. The optimization was performed using the Adam optimizer, and the loss function was defined as mean squared error (MSE) between the predicted and actual permeability residuals. We employed a single model ensemble with no dropout regularization applied during training. No additional atom or bond descriptors were provided other than the input SMILES strings.

Model performance was evaluated using root mean square error (RMSE) as the primary metric. The RMSE was calculated on the validation set at regular intervals, with model checkpoints saved after each epoch. The final model's performance on the test set was used to report the predictive accuracy of the MPNN-generated embeddings for downstream permeability prediction tasks. The final model consisted of 355,201 parameters.

**Downstream Machine Learning Modeling**

The embeddings generated from the MPNN were employed as input features for a downstream machine learning task to predict permeability residuals.

Three models were evaluated for their effectiveness at predicting permeability residuals from the MPNN-generated embeddings after hyperparameter tuning (**Fig. S6a**). All the downstream models were built within the Scikit learn framework[14] (version 1.3.2). A multilayer perceptron (MLP) demonstrated best performance on the validation set $R^2$ score, hence this model was selected for downstream use (**Fig. S6b**). While other models did achieve higher performance on the training set, their worse performance on the validation set indicates they were overfit. To verify the need for embeddings, we also trained the three models on input features generated by RDKit and by ECFP, which provide fixed representations of chemicals based on pre-determined rules. Models trained on fixed representations performed worse than embeddings-based models (**Fig. S6a**).

The MLP was configured with four hidden layers consisting of 300, 200, 32, and 16 neurons, respectively. The activation function used for all hidden layers was the ReLU function. To control overfitting, L2 regularization was applied with an alpha parameter set to 0.01. The learning rate was adaptive, with an initial value of 0.001, and the model's optimization was performed using stochastic gradient descent.

The dataset was split into training, validation, and test sets using a random 80-10-10 split. A random data split was employed in this stage in order to accurately measure the predictive performance of the model on compounds that are within the distribution of chemicals in the experimental dataset. The model was trained for a maximum of 400 epochs, with early stopping implemented to halt training when the validation loss did not improve for 10 consecutive epochs. During each epoch, the model was trained on the training set, and predictions were generated for both the training and validation sets (**Fig. S6b**). The mean squared error (MSE) was computed for both sets to monitor performance and detect the optimal stopping point.

To identify the optimal model, the epoch with the lowest validation loss was selected. This model was then re-trained from scratch for the corresponding number of epochs using the training set. The final model was evaluated on both the validation and test sets to ensure its performance generalized well to unseen data. The training and validation loss values were recorded across epochs to visualize the training process and confirm the model's convergence. Parity plots were generated to compare the predicted permeability residuals against the actual values (**Fig. 3b**). All machine plots provided in this study were made using Matplotlib (version 3.10.0).

To further interrogate the confidence of our model, we interrogated the uncertainty by performing various iterations on 30 different randomly selected model states and measuring the standard deviation of the predicted residuals for all compounds in the test set. We observed only a few points with high variance (**Fig. S6c**).

**Model Interpretability using a gradient boosted approach**

We performed an interpretability analysis to determine which features of the input data lead the model to make its predictions. A fundamental issue with the black box nature of deep learning models like the MPNN is that our embeddings don't have human-interpretable meanings. To overcome this problem, an extreme gradient boosting (XGBoost) regressor (version 2.1.3) was employed to model the relationship between interpretable molecular descriptors and the predictions made by the MycoPermeNet model[15]. The molecular descriptors used here were the 26 RDkit descriptors (RDkit version 2024.9.5), where each descriptor is an interpretable entry (e.g. molecular weight, lipophilicity of the compound, etc). Features were hand-selected features via scientific intuition for relevance to Mtb permeability, resulting in an input matrix of 1556 by 26 for our analysis.

The XGBoost model was configured with a maximum tree depth of 3 and a regularization parameter of 5 to control model complexity and prevent overfitting. The model was trained on the cleaned training set of descriptors. The trained XGBoost model was then used to predict the outcomes on the test set of interpretable descriptors. These predictions were compared against the predictions made by the original complex MLP model. The performance of the XGBoost model was evaluated by calculating the mean squared error (MSE) and the $R^2$ score between the XGBoost predictions and the complex model's predictions on the test set. These metrics provided insight into how well the simpler, interpretable model could replicate the behavior of the complex model.

To interpret the predictions made by the XGBoost model, SHAP (SHapley Additive exPlanations; version 0.5.6) was used[16]. SHAP is a method that assigns each feature an importance value based on its contribution to the model's predictions using Shapley values from cooperative game theory. SHAP values were computed to quantify the contribution of each feature (interpretable descriptor) to the predictions made by the XGBoost model.

### 3. Molecules synthesis and characterization

#### *Solid-phase synthesis of W1-3 peptides*

W1-3 peptides were prepared by standard Fmoc-based solid-phase chemistry using the rink amide resin. Briefly, to a 25 mL peptide synthesis vessel, appropriate amount of resin was added, followed by 20% piperidine in *N, N*-Dimethylformamide (DMF, 15 mL). This was followed by shaking at room temperature for 30 min. The resin was then washed with methanol ($CH_3OH$) and dichloromethane (DCM) three times. After the last wash, 4 eqiv. of amino acid was added along with 4 eqiv. of ethyl cyanohydroxyiminoacetate (Oxyma) and 4 eqiv. of *N, N'*-Diisopropylcarbodiimide (DIC). The resin was shaken at room temperature for 2 h, then washed with $CH_3OH$/DCM. The remainder of the amino acids were coupled in the same manner. Peptides were cleaved from the resin using a TFA/TIPS/$H_2O$ mixture (95:2.5:2.5, v/v/v) with shaking at room temperature for 2 h unless otherwise mentioned. The solution was filtered and concentrated prior to precipitation by the addition of cold diethyl ether to yield crude peptide. Crude peptides were purified by reverse-phased preparative high-performance liquid chromatography (RP-HPLC) equipped with Waters 1525 with 2489 UV/Visible Detector on a Phenomenex Luna 10 μm C8(2) 100 Å (250 x 21.2 mm) column using gradient elution with $H_2O$/MeOH with 0.1% TFA at 10 mL/min. The HPLC fractions of the desired purified compounds were first concentrated under reduced pressure using a rotary evaporator. The final concentrated aqueous solutions were lyophilized to dryness using Labconco Freezone 4.5 L (-84$^{o}$C) lyophilizer. The peptides were analyzed for purity using Phenomenex Luna 5 µm C8(2) with gradient elution in $H_2O$/MeCN with 0.1% TFA at 1 mL/min. Peptide identities were confirmed via high resolution electrospray ionization mass spectrometry (HRMS, ESI/MS). Analyses were obtained on an Agilent 6545B Q-TOF LC/MS equipped with 1260 infinity II LC system with auto sampler.

### *Characterization of W1-3 peptides*

### Chemical structure of W1

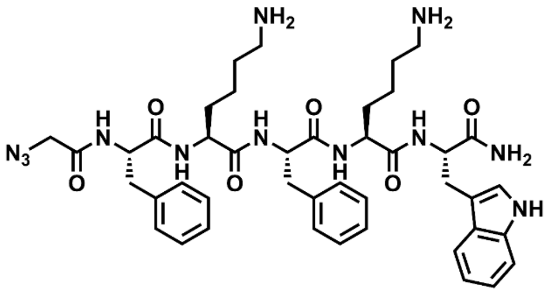


### Analytical HPLC of W1

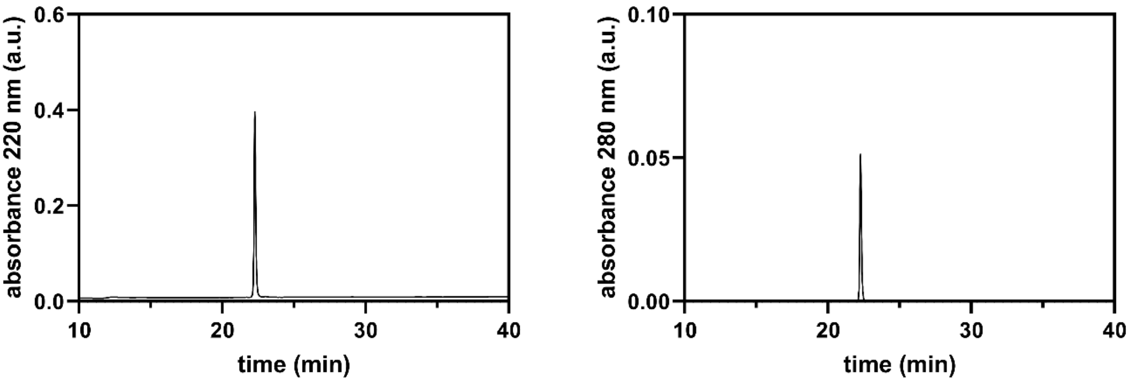


### HRMS of W1

Expected m/z: $[M+H]^+$ 837.4519; found 837.4520

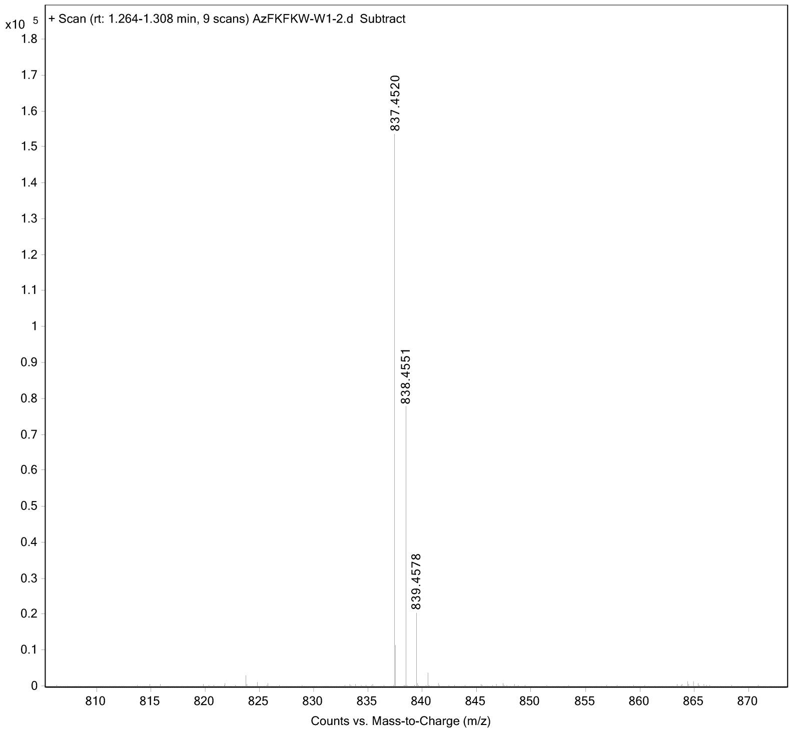

### Chemical structure of W2

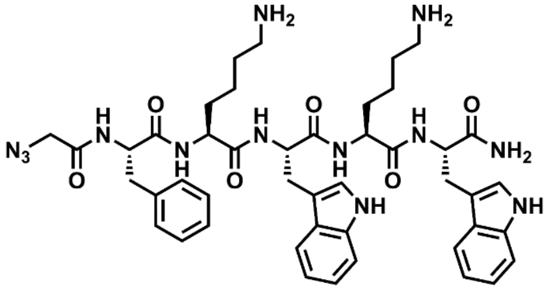


### Analytical HPLC of W2

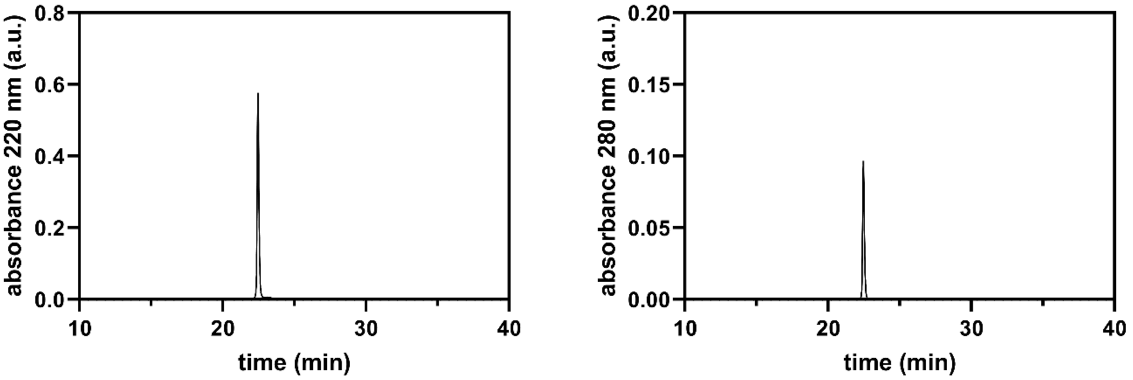


### HRMS of W2

Expected m/z: $[M+H]^+$ 876.4628; found 876.4614

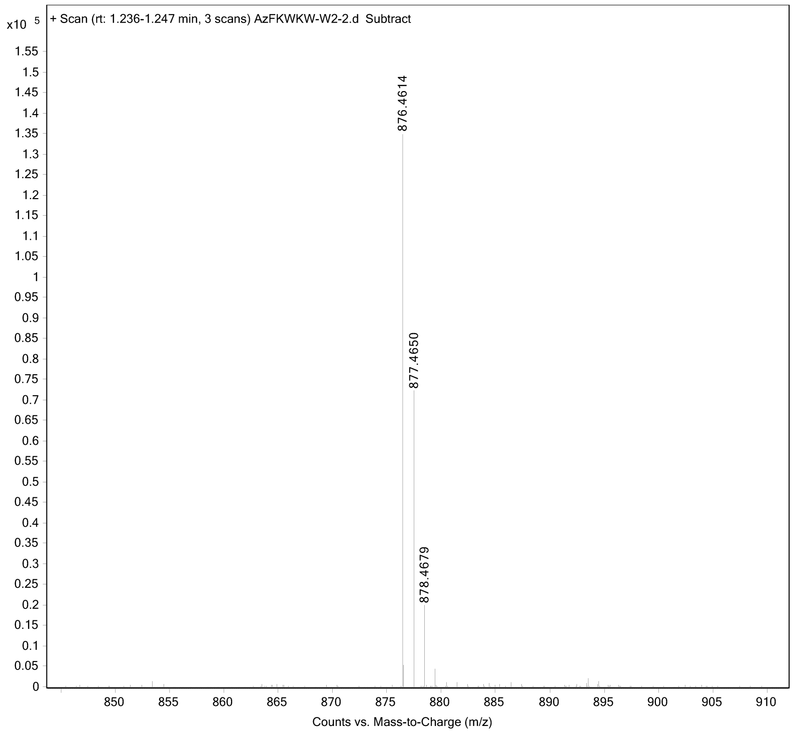

### Chemical structure of W3

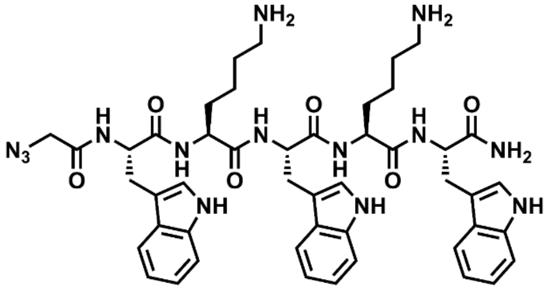


### Analytical HPLC of W3

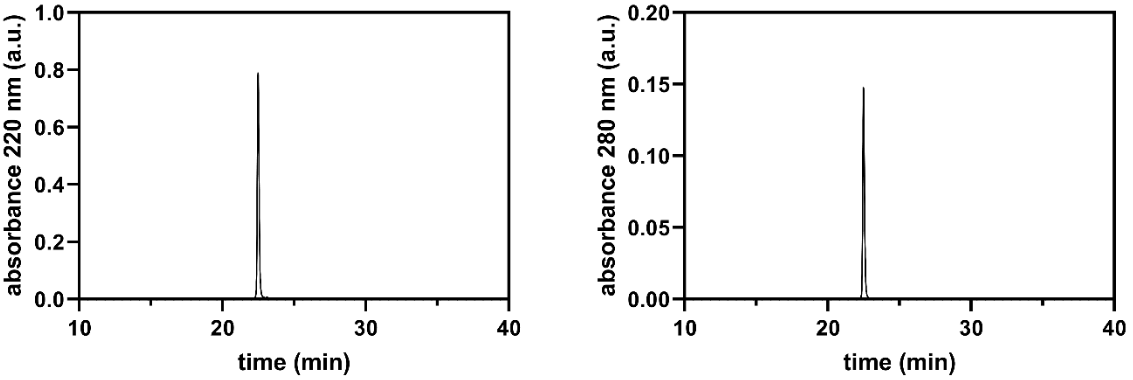


### HRMS of W3

Expected m/z: $[M+H]^+$ 915.4737; found 915.4739

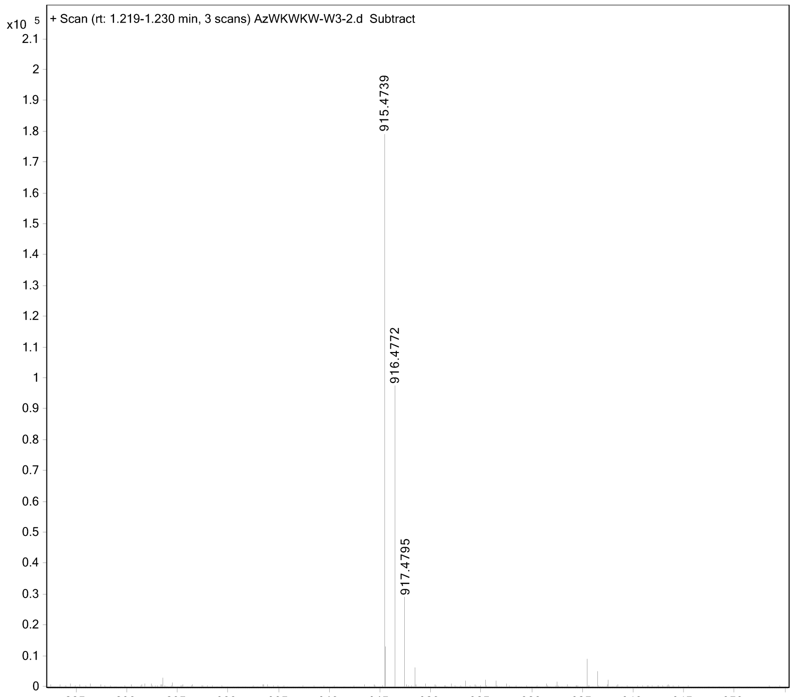

***Synthesis of JSF-2985 analogues***

All reagents were purchased from commercial suppliers and used without further purification unless noted otherwise. All chemical reactions occurring solely in an anhydrous organic solvent were executed under an inert atmosphere of nitrogen or argon unless noted otherwise. Reactions conducted at rt were usually at 21 – 24 °C. Analytical TLC was carried out with Merck silica gel 60 $F_{254}$ plates. Silica gel column chromatography utilized Teledyne Isco CombiFlash Companion or Rf+ systems. $^{1}$H NMR spectra were obtained on Bruker 500 and 600 MHz instruments and are listed in parts per million downfield from TMS. $^{1}$H NMR and $^{13}$C NMR spectra were recorded on a Bruker 500 MHz instrument. Liquid chromatography-mass spectrometry (LC-MS) was conducted on an Agilent 1260 HPLC coupled to an Agilent 6120 MS. All synthesized compounds were at least 95% pure as judged by their HPLC trace at 220 or 250 nm and were characterized by the expected parent ion/s in the MS. HRMS was executed on an Agilent 6230B Accurate Mass TOF MS.

JSF-2985 and key intermediate ethyl 5-((*tert*-butoxycarbonyl)amino)-3-methyl-1*H*-indole-2-carboxylate were synthesized as reported previously(PMID: 32197094).

**5-((4-azidobutyl)sulfonamido)-3-methyl-1*H*-indole-2-carboxamide (az-JSF-2985; JSF-6020):**

Ethyl 5-amino-3-methyl-1*H*-indole-2-carboxylate trifluoroacetic acid salt: Trifluoroacetic acid (10 mL) was added dropwise to a vigorously stirring solution of ethyl 5-((*tert*-butoxycarbonyl)amino)-3-methyl-1*H*-indole-2-carboxylate (400 mg, 1.25 mmol) in 10 mL DCM. The reaction was stirred at rt for 12 h when at that point no starting material was remaining based on LC/MS analysis. The reaction mixture was concentrated *in vacuo* to give the desired product as a white, gummy semi-solid (170 mg, 0.511 mmol, 40.7%): Calculated for $C_{12}H_{15}N_2O_2$ $[M+H]^+$ = 219.1; found 219.1.

Ethyl 5-((4-chlorobutyl)sulfonamido)-3-methyl-1*H*-indole-2-carboxylate: Ethyl 5-amino-3-methyl-1*H*-indole-2-carboxylate trifluoroacetic acid salt (170 mg, 0.511 mmol) and 4-chlorobutanesulfonyl chloride (98 mg, 0.511 mmol) were all dissolved in 5 mL pyridine at -10 °C. The reaction was stirred at -10 °C for 5 h. The reaction was diluted 10-fold with ethyl acetate and washed with 1N $HCl_{(aq)}$ solution. The organic phase was dried over anhydrous $Na_2SO_4$ and concentrated *in vacuo* to afford the desired product as a brown gummy liquid (159 mg, 0.427 mmol, 83.6%): Calculated for $C_{16}H_{22}ClN_2O_4S$ $[M+H]^+$ = 373.1; found 373.1.

Ethyl 5-((4-azidobutyl)sulfonamido)-3-methyl-1*H*-indole-2-carboxylate: To the mixture of ethyl 5-((4-chlorobutyl)sulfonamido)-3-methyl-1*H*-indole-2-carboxylate (159 mg, 0.427 mmol) in 5 mL DMSO was added sodium azide (83.0 mg, 1.28 mmol, 3 equiv). The reaction was heated to 80 °C and stirred overnight. The mixture was subjected to the addition of ice-cold water and extracted with ethyl acetate. The organic phase was dried over anhydrous $Na_2SO_4$, concentrated *in vacuo*, and purified by silica gel flash column chromatography using a gradient of 0 – 5% MeOH/DCM to afford the product as a brown solid (101 mg, 0.266 mmol, 62.3%): Calculated for $C_{16}H_{22}N_5O_4S$ $[M+H]^+$ = 380.1; found 380.1.

5-((4-azidobutyl)sulfonamido)-3-methyl-1*H*-indole-2-carboxylic acid: To a solution of ethyl 5-((4-azidobutyl)sulfonamido)-3-methyl-1*H*-indole-2-carboxylate (101 mg, 0.266 mmol) in 7 mL 1,4-dioxane was added 3 mL of an aqueous solution of LiOH (56 mg, 1.33 mmol, 5 equiv). The reaction was heated to 70 °C and stirred overnight. The mixture was cooled in an ice bath and 6N $HCl_{(aq)}$ was added slowly. The solution was extracted with ethyl acetate. The organic phase was dried over anhydrous $Na_2SO_4$ and concentrated *in vacuo* to afford a chromatographically inseparable mixture containing the desired product and two major impurities (m/z = 312.1 and m/z = 366.1) (61 mg): Calculated for $C_{14}H_{18}N_5O_4S$ $[M+H]^+$ = 352.1; found 352.0. The inseparable mixture was used in the next step without further purification.

Perfluorophenyl 5-((4-azidobutyl)sulfonamido)-3-methyl-1*H*-indole-2-carboxylate: The impure mixture containing 5-((4-azidobutyl)sulfonamido)-3-methyl-1*H*-indole-2-carboxylic acid (61 mg, ≤0.173 mmol), 1-ethyl-3-(3-dimethylaminopropyl)carbodiimide hydrochloride (50.0 mg, 0.260 mmol, ≥1.5 equiv), and pentafluorophenol (48.0 mg, 0.260 mmol, ≥1.5 equiv) were dissolved in 15 mL dichloromethane. The reaction was stirred at rt overnight. The reaction was diluted 10-fold with dichloromethane and washed with saturated $NH_4Cl_{(aq)}$ and then saturated aqueous brine solution. The organic phase was separated, dried over anhydrous $Na_2SO_4$, and concentrated *in vacuo* to afford the product as a brown solid (54 mg, 0.10 mmol, ≥60%): Calculated for $C_{20}H_{17}F_5N_5O_4S$ $[M + H]^+$ = 518.1; found 518.1.

To a solution of perfluorophenyl 5-((4-azidobutyl)sulfonamido)-3-methyl-1H-indole-2-carboxylate (54 mg, 0.104 mmol) in 5 mL acetonitrile was added ammonium hydroxide (1 mL). The reaction was stirred at rt for 12 h. After completion of the reaction as determined by TLC and LC/MS, the crude reaction mixture was taken up into EtOAc and washed with saturated $NH_4Cl_{(aq)}$ solution and a saturated aqueous brine solution. The organic phase was dried over anhydrous $Na_2SO_4$ and purified by reverse phase column chromatography using a gradient of 0 – 60% $H_2O$/ACN to afford the title product as a white solid (16 mg, 0.044 mmol, 42% over 2 steps): $^1$H NMR (500 MHz, $d_6$-DMSO) δ 11.2 (s, 1H), 9.42 (s, 1H), 7.37 (m, 4H), 7.11 (dd, $J$ = 8.7, 2.0 Hz, 1H), 3.31 (s, 2H), 3.00 (m, 2H), 2.45 (s, 3H), 1.75 (p, $J$ = 7.6 Hz, 2H), 1.59 (p, $J$ = 7.0 Hz, 2H). Also noted, 3.3 (s, $H_2O$). Calculated for $C_{14}H_{19}N_6O_3S$ $[M+H]^+$ = 351.1239; found 351.1238.

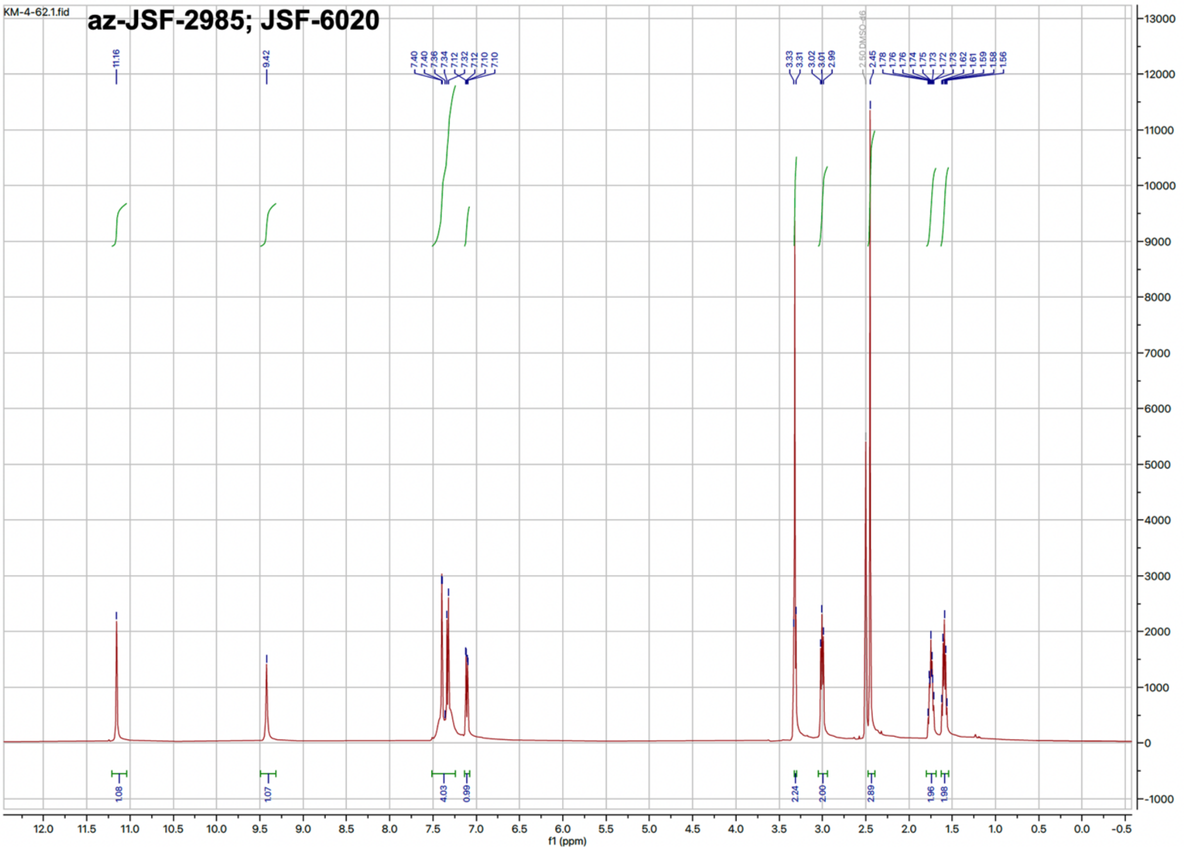


**N-(2-(1H-imidazol-2-yl)ethyl)-5-((4-azidobutyl)sulfonamido)-3-methyl-1H-indole-2-carboxamide (az-JSF-2985-imidazole; JSF-5098):**

Perfluorophenyl 5-((*tert*-butoxycarbonyl)amino)-3-methyl-1*H*-indole-2-carboxylate: 5-((*tert*-butoxycarbonyl)amino)-3-methyl-1*H*-indole-2-carboxylic acid (300 mg, 1.03 mmol), 1-ethyl-3-(3-dimethylaminopropyl)carbodiimide hydrochloride (296 mg, 1.55 mmol, 1.5 equiv), and pentafluorophenol (276 mg, 1.55 mmol, 1.5 equiv) were dissolved in 30 mL dichloromethane. The reaction was stirred at rt overnight. The reaction was diluted 10-fold with dichloromethane and washed with saturated $NH_4Cl_{(aq)}$ and then saturated aqueous brine solution. The organic phase was separated, dried over anhydrous $Na_2SO_4$, and concentrated *in vacuo* to afford the product as an off-white solid (318 mg, 0.697 mmol, 67.5%): Calculated for $C_{21}H_{18}F_5N_2O_4$ $[M + H - 56]^+$ = 401.1; found 401.0.

*tert*-butyl (2-((2-(1*H*-imidazol-2-yl)ethyl)carbamoyl)-3-methyl-1*H*-indol-5-yl)carbamate: To the pentafluoro ester (180 mg, 0.394 mmol) in 10 mL acetonitrile was added 2-(1*H*-imidazol-2-yl)ethan-1-amine hydrochloride (218 mg, 1.18 mmol, 3 equiv). The reaction was stirred at rt for 4 h. After completion of the reaction as determined by TLC and LC/MS, the crude reaction mixture was taken up into EtOAc and washed with saturated $NH_4Cl_{(aq)}$ solution and then saturated aqueous brine solution. The organic phase was dried over anhydrous $Na_2SO_4$ and purified by silica gel flash column chromatography using a gradient of 0 – 20% MeOH/DCM to afford the desired product as a white solid (83 mg, 0.21 mmol, 55%): Calculated for $C_{20}H_{26}N_5O_3$ $[M+H]^+$ = 384.2; found 384.2.

*N*-(2-(1*H*-imidazol-2-yl)ethyl)-5-amino-3-methyl-1*H*-indole-2-carboxamide trifluoroacetic acid salt: Trifluoroacetic acid (5 mL) was added dropwise to a vigorously stirring solution of *tert*-butyl (2-((2-(1*H*-imidazol-2-yl)ethyl)carbamoyl)-3-methyl-1*H*-indol-5-yl)carbamate (150 mg, 0.391 mmol) in 15 mL DCM. The reaction was stirred at rt for 12 h when no starting material was remaining based on LC/MS analysis. The reaction mixture was concentrated *in vacuo* to give the desired product as a white solid (50 mg, 0.13 mmol, 33%): Calculated for $C_{15}H_{18}N_5O$ $[M+H]^+$ = 284.1; found 284.1.

*N*-(2-(1*H*-imidazol-2-yl)ethyl)-5-((4-chlorobutyl)sulfonamido)-3-methyl-1*H*-indole-2-carboxamide: *N*-(2-(1*H*-imidazol-2-yl)ethyl)-5-amino-3-methyl-1*H*-indole-2-carboxamide trifluoroacetic acid salt (50 mg, 0.13 mmol), and 4-chlorobutanesulfonyl chloride (37 mg, 0.19 mmol, 1.5 equiv) were all dissolved in 5 mL pyridine at 0 °C. The reaction was stirred at rt for 5 h. The reaction was diluted 10-fold with ethyl acetate and washed with with 1N $HCl_{(aq)}$ solution. The organic phase was dried over anhydrous $Na_2SO_4$ and concentrated *in vacuo* to afford an inseparable mixture containing the desired product (43 mg): Calculated for $C_{19}H_{25}ClN_5O_3S$ $[M+H]^+$ = 438.1; found 438.1. The inseparable mixture was used in the next step without further purification.

To the impure mixture containing 5-((4-chlorobutyl)sulfonamido)-*N*-(2-cyclopentylethyl)-3-methyl-1*H*-indole-2-carboxamide (40 mg) in 5 mL DMSO was added sodium azide (18.0 mg, 0.274 mmol). The reaction was heated to 80 °C and stirred overnight. The mixture was subjected to the addition of ice-cold water and extracted with ethyl acetate. The organic phase was dried over anhydrous $Na_2SO_4$, concentrated *in vacuo*, and purified by silica gel flash column chromatography using a gradient of 0 – 30% EtOAc/hexanes and then 0 – 5% MeOH/DCM to afford the title product as an off-white solid (5.0 mg, 0.011 mmol, 8.4% based on two steps): $^1$H NMR (500 MHz, MeOD) δ 7.50 (s, 1H), 7.35 (d, *J* = 8.7 Hz, 1H), 7.18 (dd, *J* = 8.7, 2.1 Hz, 1H), 6.97 (s, 2H), 3.75 (t, *J* = 7.0 Hz, 2H), 3.29 (d, *J* = 6.6 Hz, 2H), 3.04 (m, 4H), 2.46 (s, 3H), 1.98 (m, 2H), 1.66 (p, *J* = 6.9 Hz, 2H): Also, noted 3.3 (s, $H_2O$) and 1.3 (s). 4 NHs were unaccounted for, presumably due to H/D exchange with the NMR solvent. Calculated for $C_{19}H_{25}N_8O_3S$ $[M+H]^+$ = 445.1770; found 445.1754.

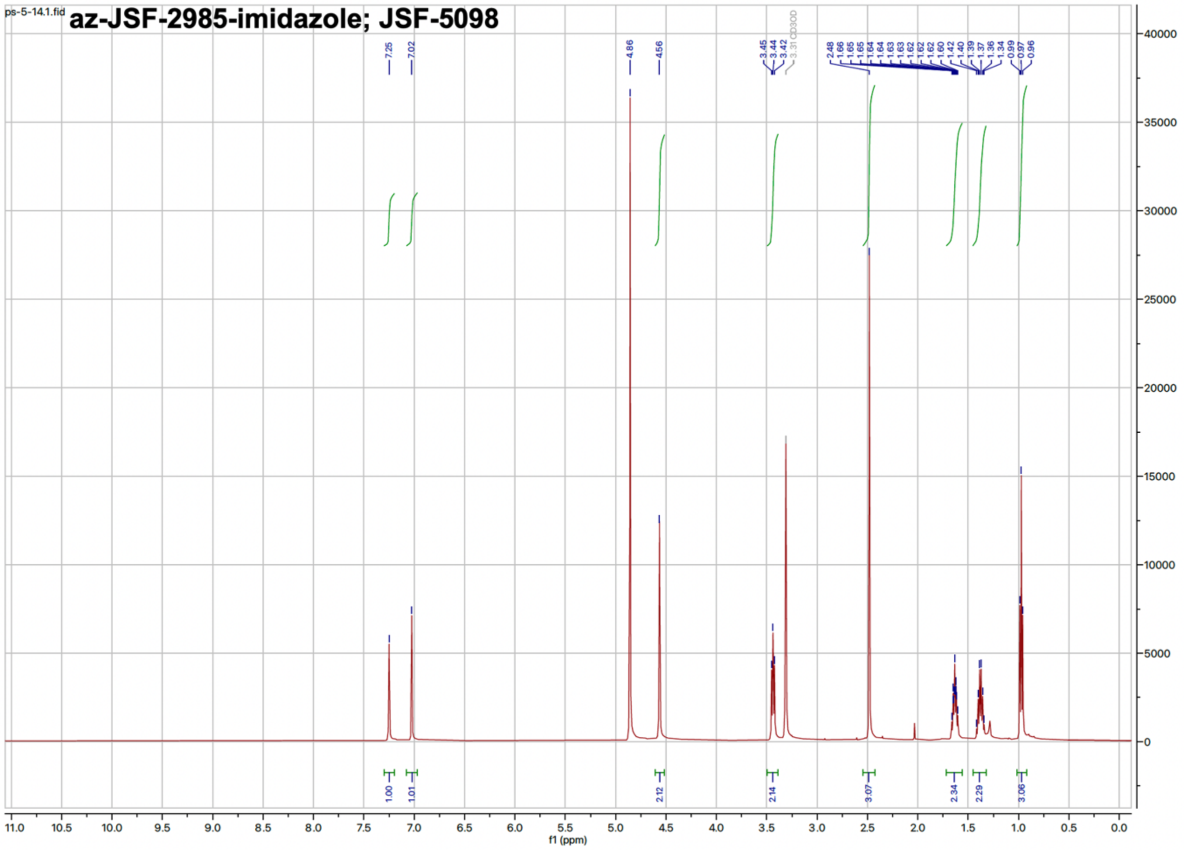


***N*-(2-(1*H*-pyrazol-3-yl)ethyl)-5-((4-azidobutyl)sulfonamido)-3-methyl-1*H*-indole-2-carboxamide (az-JSF-2985-pyrazole; JSF-6009):**

Perfluorophenyl 5-((*tert*-butoxycarbonyl)amino)-3-methyl-1*H*-indole-2-carboxylate: 5-((*tert*-butoxycarbonyl)amino)-3-methyl-1*H*-indole-2-carboxylic acid (207 mg, 0.713 mmol), 1-ethyl-3-(3-dimethylaminopropyl)carbodiimide hydrochloride (204 mg, 1.07

mmol, 1.5 equiv), and pentafluorophenol (197 mg, 1.07 mmol, 1.5 equiv) were dissolved in 30 mL dichloromethane. The reaction was stirred at rt overnight. The reaction was diluted 10-fold with dichloromethane and washed with saturated $NH_4Cl_{(aq)}$ and followed by a wash with saturated aqueous brine solution. The organic phase was separated, dried over anhydrous $Na_2SO_4$, and concentrated *in vacuo* to afford the desired product as an off-white solid (290 mg, 0.635 mmol, 89.1%): Calculated for $C_{21}H_{18}F_5N_2O_4$ $[M + H - 56]^+$ = 401.1; found 401.0.

*tert*-butyl (2-((2-(1*H*-pyrazol-3-yl)ethyl)carbamoyl)-3-methyl-1*H*-indol-5-yl)carbamate: To the pentafluoro ester (290 mg, 0.635 mmol) in 10 mL acetonitrile was added 2-(1*H*-pyrazol-3-yl)ethan-1-amine hydrochloride (234 mg, 1.27 mmol, 3 equiv). The reaction was stirred at rt for 4 h. After completion of the reaction as determined by TLC and LC/MS, the crude reaction mixture was taken up into EtOAc and washed with saturated $NH_4Cl_{(aq)}$ solution and a saturated aqueous brine solution. The organic phase was dried over anhydrous $Na_2SO_4$ and purified by silica gel flash column chromatography using a gradient of 0 – 10% MeOH/DCM to afford the product as a yellow solid (210 mg, 0.548 mmol, 86.4%): Calculated for $C_{20}H_{26}N_5O_3$ $[M+H]^+$ = 384.2; found 384.2.

*N*-(2-(1*H*-pyrazol-3-yl)ethyl)-5-amino-3-methyl-1*H*-indole-2-carboxamide trifluoroacetic acid salt: Trifluoroacetic acid (5 mL) was added dropwise to a vigorously stirring solution of *tert*-butyl (2-((2-(1*H*-pyrazol-3-yl)ethyl)carbamoyl)-3-methyl-1*H*-indol-5-yl)carbamate (210 mg, 0.548 mmol) in 10 mL DCM. The reaction was stirred at rt for 12 h after which no starting material was remaining based on LC/MS analysis. The reaction mixture was concentrated *in vacuo* to give the desired product as a brown, gummy semi-solid (105 mg, 0.371 mmol, 67.7%): Calculated for $C_{15}H_{18}N_5O$ $[M+H]^+$ = 284.2; found 284.1.

*N*-(2-(1*H*-pyrazol-3-yl)ethyl)-5-((4-chlorobutyl)sulfonamido)-3-methyl-1*H*-indole-2-carboxamide: *N*-(2-(1*H*-pyrazol-3-yl)ethyl)-5-amino-3-methyl-1*H*-indole-2-carboxamide trifluoroacetic acid salt (105 mg, 0.371 mmol) and 4-chlorobutanesulfonyl chloride (78 mg, 0.41 mmol) were dissolved in 5 mL pyridine at 0 °C. The reaction was stirred at rt for 5 h. The reaction was diluted 10-fold with ethyl acetate and washed with 1N $HCl_{(aq)}$ solution. The organic phase was dried over anhydrous $Na_2SO_4$ and concentrated *in vacuo* to afford a chromatographically inseparable mixture containing the desired product and two major impurities (m/z =

534.1 and m/z = 380.1) (59 mg): Calculated for $C_{19}H_{25}ClN_5O_3S$ $[M+H]^+$ = 438.1; found 438.1. The inseparable mixture was used in the next step without further purification.

To the mixture containing *N*-(2-(1*H*-pyrazol-3-yl)ethyl)-5-((4-chlorobutyl)sulfonamido)-3-methyl-1*H*-indole-2-carboxamide (59 mg) in 5 mL DMSO was added sodium azide (26.0 mg, 0.405 mmol). The reaction was heated to 80 °C and stirred overnight. The mixture was subjected to the addition of ice-cold water and extracted with ethyl acetate. The organic phase was dried over anhydrous $Na_2SO_4$, concentrated *in vacuo*, and purified by silica gel flash column chromatography using a gradient of 0 – 30% EtOAc/hexanes and then 0 – 5% MeOH/DCM to afford the title product as a white solid (2.2 mg, 0.0049 mmol, 1.2% based on two steps): $^1$H NMR (500 MHz, $d_6$-DMSO) δ 12.5 (br s, 1H), 11.2 (s, 1H), 7.90 (d, *J* = 14.8 Hz, 2H), 7.33 (m, 3H), 7.09 (dd, *J* = 8.7, 2.1 Hz, 1H), 6.12 (s, 1H), 3.55 (q, *J* = 6.8 Hz, 2H), 2.99 (m, 2H), 2.87 (s, 2H), 2.42 (s, 3H), 1.74 (m, 2H), 1.59 (p, *J* = 7.0 Hz, 2H). 1 NH was unaccounted for, presumably due to H/D exchange with the NMR solvent. Also noted 1.2 (s). Calculated for $C_{19}H_{25}N_8O_3S$ $[M+H]^+$ = 445.1770; found 445.1768.

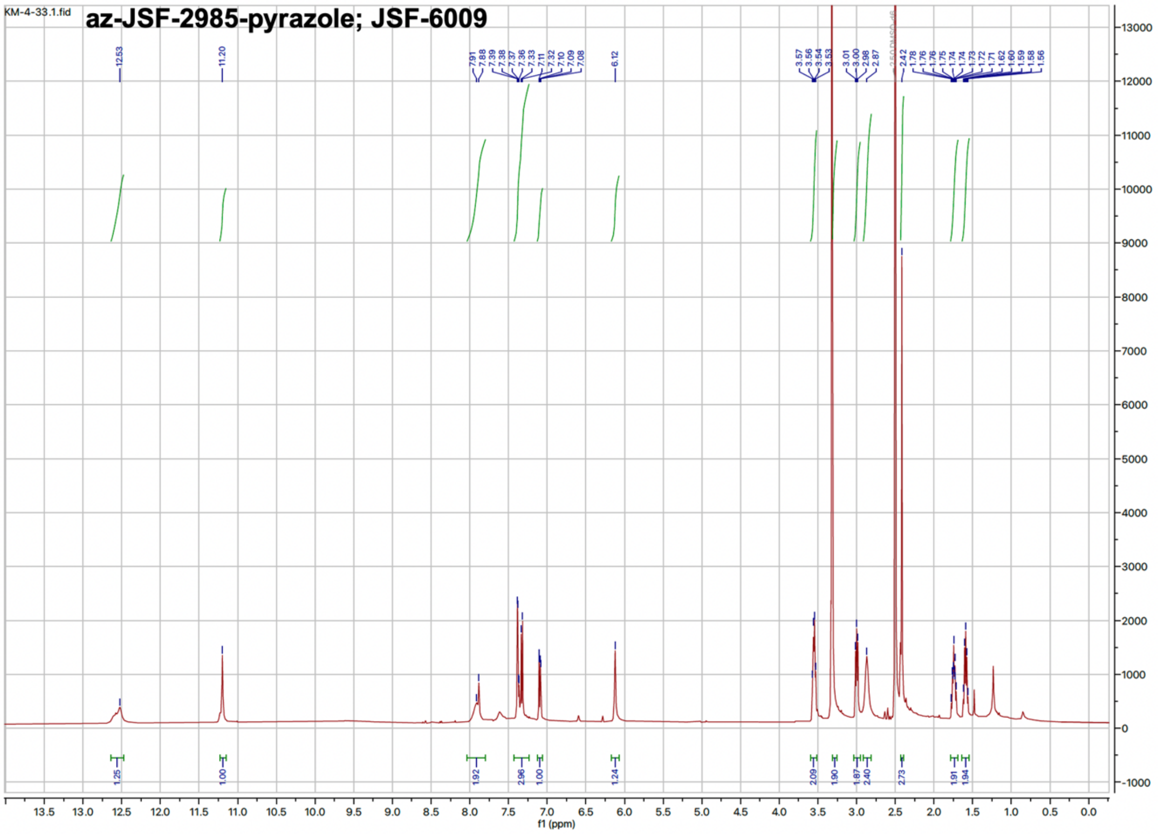

**5-((4-azidobutyl)sulfonamido)-3-methyl-*N*-(2-(pyrrolidin-3-yl)ethyl)-1*H*-indole-2-carboxamide (az-JSF-2985-pyrrolidine; JSF-6001):**

Perfluorophenyl 5-((*tert*-butoxycarbonyl)amino)-3-methyl-1*H*-indole-2-carboxylate: 5-((*tert*-butoxycarbonyl)amino)-3-methyl-1*H*-indole-2-carboxylic acid (238 mg, 0.820 mmol, 1.5 equiv), 1-ethyl-3-(3-dimethylaminopropyl)carbodiimide hydrochloride (235 mg, 1.23 mmol, 1.5 equiv), and pentafluorophenol (226 mg, 1.23 mmol, 1.5 equiv) were dissolved in 20 mL dichloromethane. The reaction was stirred at rt overnight. The reaction was diluted 10-fold with dichloromethane and washed with saturated $NH_4Cl_{(aq)}$ and then saturated aqueous brine solution. The organic phase was separated, dried over anhydrous $Na_2SO_4$, and concentrated *in vacuo* to afford the product as a brown solid (270 mg, 0.592 mmol, 72.2%): Calculated for $C_{21}H_{18}F_5N_2O_4$ $[M + H -56]^+$ = 401.0; found 401.0.

Perfluorophenyl 5-amino-3-methyl-1*H*-indole-2-carboxylate: Trifluoroacetic acid (3 mL) was added dropwise to a vigorously stirring solution of perfluorophenyl 5-((*tert*-butoxycarbonyl)amino)-3-methyl-1*H*-indole-2-carboxylate (200 mg, 0.438 mmol) in 15 mL DCM. The reaction was stirred at rt for 3 h when no starting material was remaining based on LC/MS analysis. The reaction was neutralized with saturated aqueous $NaHCO_3$ solution until the pH of the aqueous layer was approximately 8. The DCM layer was collected, and the remaining aqueous phase was extracted twice with EtOAc. The organic fractions were pooled and washed with saturated aqueous brine solution. The organic phase was dried over anhydrous $Na_2SO_4$ and concentrated *in vacuo* to give the desired product as a brown solid (117 mg, 0.328 mmol, 74.9%): Calculated for $C_{16}H_{10}F_5N_2O_2$ $[M+H]^+$ = 357.1; found 357.0.

*tert*-butyl 3-(2-(5-amino-3-methyl-1*H*-indole-2-carboxamido)ethyl)pyrrolidine-1-carboxylate: To a solution of perfluorophenyl 5-amino-3-methyl-1*H*-indole-2-carboxylate (117 mg, 0.328 mmol) in 10 mL acetonitrile was added *tert*-butyl 3-(2-aminoethyl)pyrrolidine-1-carboxylate (141 mg, 0.657 mmol, 2 equiv). The reaction was stirred at rt for 12 h. After completion of reaction as determined by TLC and LC/MS, the reaction mixture was taken up into EtOAc and washed with saturated $NH_4Cl_{(aq)}$ solution and a saturated aqueous brine solution. The organic phase was dried over anhydrous $Na_2SO_4$ and purified by silica gel flash column chromatography using a gradient of 0 – 10% MeOH/DCM to afford the desired product as a white solid (83.0 mg, 0.215 mmol, 65.3%): Calculated for $C_{21}H_{31}N_4O_3$ $[M+H-56]^+$ = 331.2; found 331.1.

*tert*-butyl 3-(2-(5-((4-chlorobutyl)sulfonamido)-3-methyl-1*H*-indole-2-carboxamido)ethyl) pyrrolidine-1-carboxylate: *tert*-butyl 3-(2-(5-amino-3-methyl-1*H*-indole-2-carboxamido)ethyl)pyrrolidine-1-carboxylate (83.0 mg, 0.215 mmol), and 4-chlorobutanesulfonyl chloride (45.0 mg, 0.236 mmol) were dissolved in 5 mL pyridine at 0 °C. The reaction was stirred for 4 h at 0 °C. After completion of the reaction as determined by TLC and LC/MS, the crude residue was diluted 10-fold with ethyl acetate and washed with with 1N $HCl_{(aq)}$ solution. The organic phase was dried over anhydrous $Na_2SO_4$, concentrated *in vacuo*, and purified by silica gel flash column chromatography using a gradient of 0 – 10% MeOH/DCM to afford the desired product as a white gummy liquid (65.0 mg, 0.120 mmol, 56.0%): Calculated for $C_{25}H_{38}ClN_4O_5S$ $[M+H-100]^+$ = 441.2; found 441.1.

5-((4-chlorobutyl)sulfonamido)-3-methyl-*N*-(2-(pyrrolidin-3-yl)ethyl)-1*H*-indole-2-carboxamide: Trifluoroacetic acid (2 mL) was added dropwise to a vigorously stirring solution of *tert*-butyl 3-(2-(5-((4-chlorobutyl)sulfonamido)-3-methyl-1*H*-indole-2-carboxamido)ethyl) pyrrolidine-1-carboxylate (65.0 mg, 0.120 mmol) in 10 mL DCM. The reaction was stirred at rt for 5 h when no starting material was remaining based on LC/MS analysis. The reaction was neutralized with saturated aqueous $NaHCO_3$ solution until the pH of the aqueous layer was approximately 8. The DCM layer was collected, and the remaining aqueous phase was extracted twice with 5% MeOH/DCM. The organic fractions were pooled and washed with saturated aqueous brine solution. The organic phase was dried over anhydrous $Na_2SO_4$ and concentrated *in vacuo* to give the desired product as a white gummy semi-solid (43.0 mg, 0.097 mmol, 81.1%): Calculated for $C_{20}H_{30}ClN_4O_3S$ $[M+H]^+$ = 441.2; found 441.1.

To a solution of 5-((4-chlorobutyl)sulfonamido)-3-methyl-*N*-(2-(pyrrolidin-3-yl)ethyl)-1*H*-indole-2-carboxamide (43.0 mg, 0.097 mmol) in 3 mL DMSO was added sodium azide (19.0 mg, 0.293 mmol, 3 equiv). The reaction was heated to 80 °C and stirred overnight. The mixture was cooled with the addition of ice-cold water and extracted with ethyl acetate followed by 5% MeOH/DCM. The organic phase was dried over anhydrous $Na_2SO_4$, concentrated *in vacuo*, and purified by reverse phase column chromatography using a gradient of 0 – 100% $CH_3CN/H_2O$ to afford the title product as an off-white solid (15 mg, 0.033 mmol, 34%): $^1H$ NMR (500 MHz, $d_6$-DMSO) δ 9.28 (br s, 1H), 8.59 (s, 1H), 7.39 (s, 1H), 7.30 (d, *J* = 8.5 Hz, 1H), 7.11 (d, *J* = 6.7 Hz, 1H), 3.30 (s, 2H), 3.20 (d, *J* = 11.4 Hz, 2H), 3.00 (t, *J* = 7.8 Hz, 3H), 2.70 (t, *J* = 9.9 Hz, 1H), 2.46 (s, 3H), 2.28 (m, 1H), 2.12 (d, *J* = 7.8 Hz, 1H), 2.0 (m, 1H), 1.73 (dt, *J* = 15.0, 7.6

Hz, 3H), 1.66 (dd, $J$ = 14.7, 7.2 Hz, 2H), 1.59 (t, $J$ = 7.3 Hz, 2H), 1.49 (d, $J$ = 10.1 Hz, 2H): Also noted, 3.3 (s, $H_2O$) and 1.3 (s). One H was unaccounted for and presumably was an N*H*. Calculated for $C_{20}H_{30}N_7O_3S$ $[M+H]^+$ = 448.2131; found 448.2124.

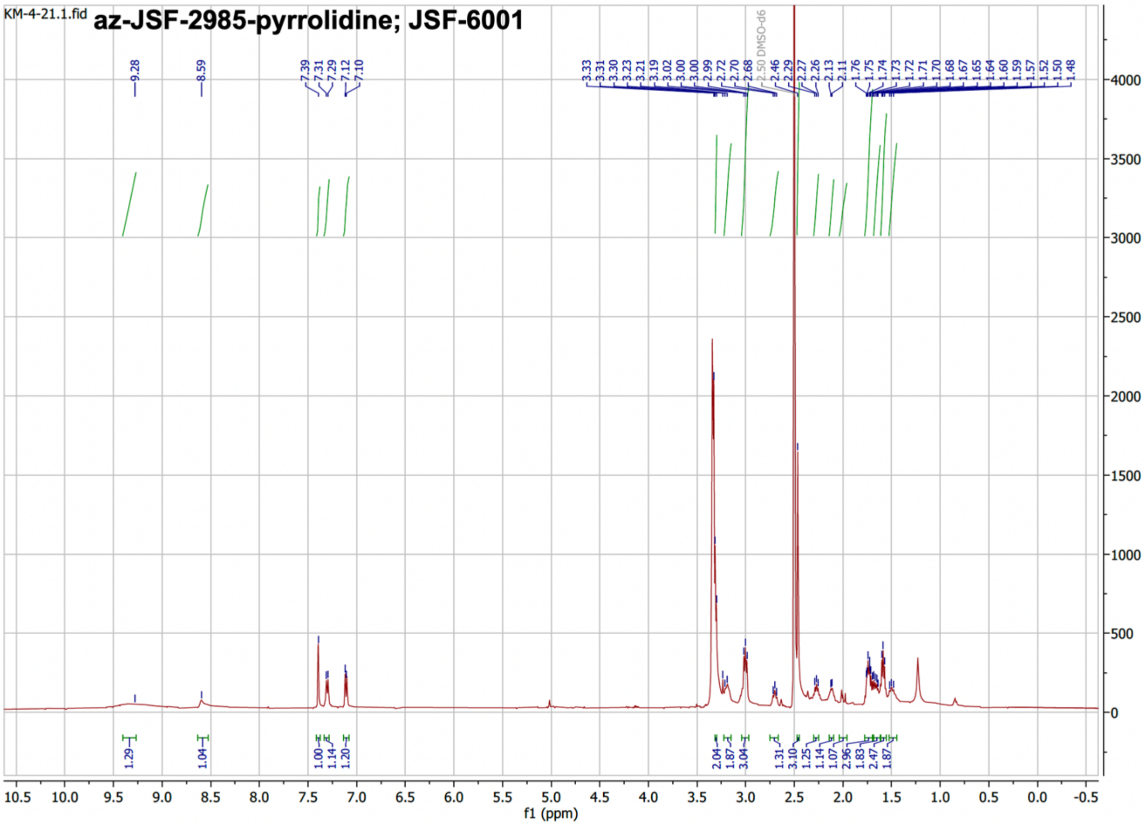


**5-((4-azidobutyl)sulfonamido)-*N*-(2-cyclopentylethyl)-3-methyl-1*H*-indole-2-carboxamide (az-JSF-2985-cyclopentane; JSF-5093):**

Perfluorophenyl 5-((*tert*-butoxycarbonyl)amino)-3-methyl-1*H*-indole-2-carboxylate: 5-((*tert*-butoxycarbonyl)amino)-3-methyl-1*H*-indole-2-carboxylic acid (400 mg, 1.37 mmol), 1-ethyl-3-(3-dimethylaminopropyl)carbodiimide hydrochloride (395 mg, 2.06 mmol), and pentafluorophenol (379 mg, 2.06 mmol) were dissolved in 30 mL dichloromethane. The reaction was stirred at rt overnight. The reaction was diluted 10-fold with dichloromethane and washed with saturated $NH_4Cl_{(aq)}$ and then saturated aqueous brine solution. The organic phase was separated, dried over anhydrous $Na_2SO_4$, and concentrated *in vacuo* to afford the desired product as an off-white solid (310 mg, 0.679 mmol, 49.3%): Calculated for $C_{21}H_{18}F_5N_2O_4$ $[M + H -56]^+$ = 401.0; found 401.0.

*tert*-butyl (2-((2-cyclopentylethyl)carbamoyl)-3-methyl-1*H*-indol-5-yl)carbamate: To the pentafluoro ester (310 mg, 0.679 mmol) in 10 mL acetonitrile was added 2-cyclopentylethan-1-amine (153 mg, 1.35 mmol, 2 equiv). The reaction was stirred at rt for 4 h. After completion of reaction as determined by TLC and LC/MS, the crude reaction mixture was taken up into EtOAc and washed with saturated $NH_4Cl_{(aq)}$ solution and then a saturated aqueous brine solution. The organic phase was dried over anhydrous $Na_2SO_4$ and purified by silica gel flash column chromatography using a gradient of 0 – 10% MeOH/DCM to afford the desired product as a yellow solid (209 mg, 0.542 mmol, 80.0%): Calculated for $C_{22}H_{32}N_3O_3$ $[M+H]^+$ = 386.2; found 386.2.

5-amino-*N*-(2-cyclopentylethyl)-3-methyl-1*H*-indole-2-carboxamide: Trifluoroacetic acid (3 mL) was added dropwise to a vigorously stirring solution of *tert*-butyl (2-((2-cyclopentylethyl) carbamoyl)-3-methyl-1*H*-indol-5-yl)carbamate (300 mg, 1.05 mmol) in 15 mL DCM. The reaction was stirred at rt for 12 h after which no starting material was remaining based on LC/MS analysis. The reaction was neutralized with saturated aqueous $NaHCO_3$ solution until the pH of the aqueous layer was approximately 8. The DCM layer was collected, and the remaining aqueous phase was extracted twice with EtOAc. The organic fractions were pooled and washed with saturated aqueous brine solution. The organic phase was dried over anhydrous $Na_2SO_4$ and concentrated *in vacuo* to give the desired product as a brown solid (210 mg, 0.736 mmol, 94.5%): Calculated for $C_{17}H_{24}N_3O$ $[M+H]^+$ = 286.2; found 286.1.

5-((4-chlorobutyl)sulfonamido)-*N*-(2-cyclopentylethyl)-3-methyl-1*H*-indole-2-carboxamide: 5-amino-*N*-(2-cyclopentylethyl)-3-methyl-1*H*-indole-2-carboxamide (210 mg, 0.736 mmol), and 4-chlorobutanesulfonyl chloride (155 mg, 0.810 mmol,1.1 equiv) were dissolved in 10 mL pyridine at 0 °C. The reaction was stirred at rt overnight. The reaction was diluted 10-fold with ethyl acetate and washed with 1N $HCl_{(aq)}$ solution. The organic phase was dried over anhydrous $Na_2SO_4$, concentrated *in vacuo*, and purified by silica gel flash column chromatography using a gradient of 0 – 5% MeOH/DCM to afford the desired product as a yellow solid (136 mg, 0.309 mmol, 42.1%): Calculated for $C_{21}H_{31}ClN_3O_3S$ $[M+H]^+$ = 440.2; found 440.1.

To a solution of 5-((4-chlorobutyl)sulfonamido)-*N*-(2-cyclopentylethyl)-3-methyl-1*H*-indole-2-carboxamide (125 mg, 0.284 mmol) in 10 mL DMSO was added sodium azide (46.0 mg, 0.711 mmol). The reaction was heated to 80 °C and stirred overnight. The mixture was subjected to the addition of ice-cold water and extracted with ethyl acetate. The organic phase was dried over anhydrous $Na_2SO_4$, concentrated *in vacuo*, and purified by silica gel flash column chromatography using a gradient of 0 – 30% EtOAc/hexanes to afford the title product as a white solid (21 mg, 0.047 mmol, 16%): $^1$H NMR (500 MHz, $d_6$-DMSO) δ 11.1 (s, 1H), 9.43 (s, 1H), 7.81 (t, *J* = 5.8 Hz, 1H), 7.39 (s, 1H), 7.33 (d, *J* = 8.7 Hz, 1H), 7.10 (dd, *J* = 8.6, 2.1 Hz, 1H), 3.30 (d, *J* = 8.9 Hz, 2H), 3.00 (m, 2H), 2.43 (s, 3H), 1.76 (dtd, *J* = 18.0, 9.7, 5.9 Hz, 5H), 1.57 (m, 6H), 1.49 (dd, *J* = 7.5, 4.7 Hz, 3H), 1.12 (q, *J* = 8.6 Hz, 3H): Also noted, 3.3 (s, $H_2O$) and 1.3 (s). Calculated for $C_{21}H_{31}N_6O_3S$ $[M+H]^+$ = 447.2178; found 447.2156.

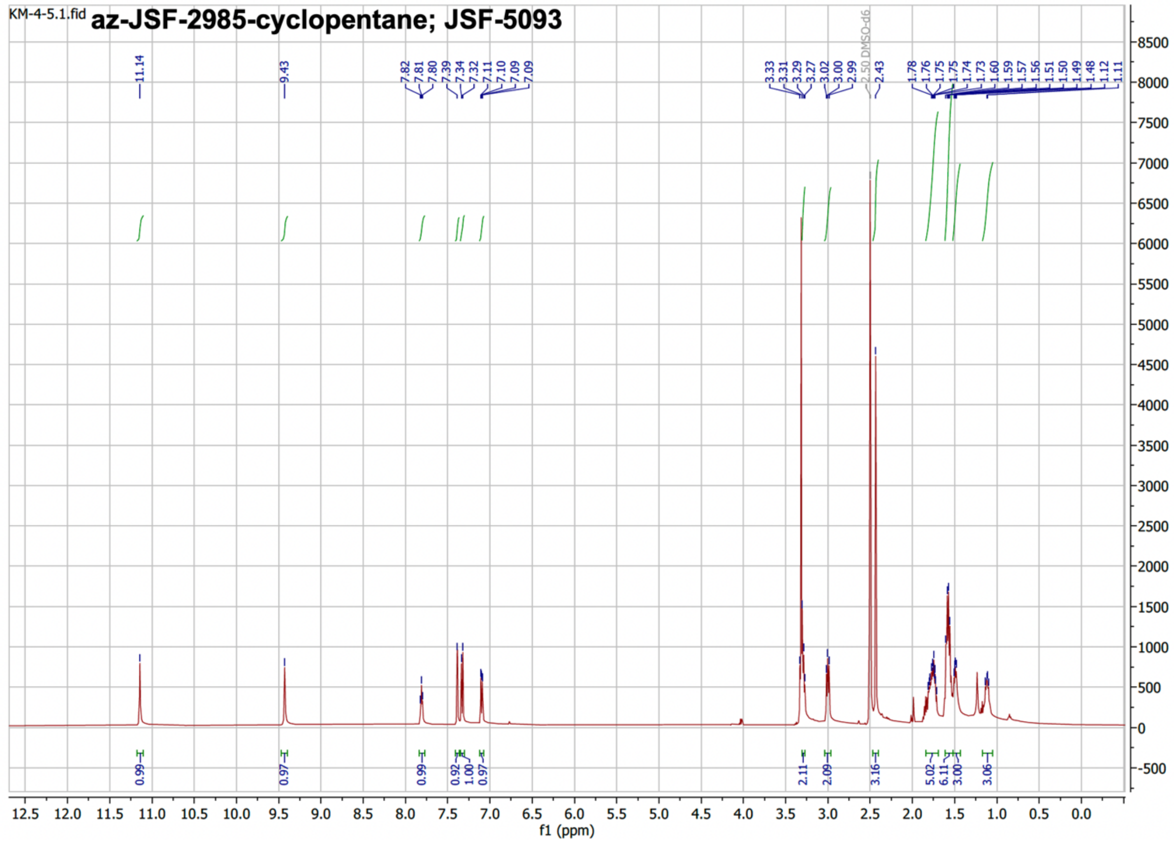
